# Dynamic Modulation of Neural Cell Identity Maintenance via Lysine Acetyltransferase Type 3 Protein Targeted Degradation

**DOI:** 10.64898/2026.10.02.756163

**Authors:** Juan Zaragoza-Lillo, Sergio Niñerola, Verónica Estefania Andrade-Montero, Francisco José López-Murcia, Tania López-Hernández, Angel Barco, Rafael Alcalá-Vida

**Affiliations:** Instituto de Neurociencias de Alicante (Universidad Miguel Hernández - Consejo Superior de Investigaciones Científicas). Avenida Santiago Ramón y Cajal, s/n, Sant Joan d’Alacant, 03550, Alicante, Spain; Department of Pathophysiology of Nervous System Diseases, Institute of Neurosciences-Maria de Maeztu Unit of Excellence, University of Barcelona, and Bellvitge Biomedical Research Institute (IDIBELL), L’Hospitalet de Llobregat, 08907 Barcelona, Spain; Department of Pathology and Experimental Therapy, Institute of Neurosciences-Maria de Maeztu Unit of Excellence, University of Barcelona, and Bellvitge Biomedical Research Institute (IDIBELL), L’Hospitalet de Llobregat, Barcelona 08907, Spain

**Keywords:** neural identity maintenance, KAT3, histone acetyltransferases, transcriptomics, PROTAC, CBP, p300, neural lineage

## Abstract

Cell identity acquisition during central nervous system development leads to the establishment of cell-specific transcriptional programs. Once acquired, this identity must be actively maintained, among other molecular mechanisms, by the acetylation of cell-type-specific regulatory regions carried out by the lysine acetyltransferases type 3 (KAT3) proteins CBP and p300. Here, we characterize the rapid transcriptional and physiological consequences of KAT3 elimination in neuronal and glial lineages by means of a KAT3-specific PROTAC degrader, providing strong evidence for their central role as neural identity wardens. Importantly, the restoration of KAT3 proteins in glial cells after cell identity erasure resulted in the recovery of the original transcriptome and physiological functions, probing the dynamic nature of neural cell identities. These findings expand the role of CBP/p300 as identity wardens and challenge our perception of terminal cellular identities as static irrevocable states, opening new avenues for neural identity manipulation with biomedical purposes.

## Introduction

Cellular specification during central nervous system (CNS) development is achieved by the orchestrated action of distinct batteries of transcription factors, tightly regulated by epigenetic regulatory mechanisms engaged at early embryonic stages (B. Yao et al., 2016). In the case of neurons this represents an additional caveat, as neuronal identity has not only to be stablished, but needs to be actively maintained throughout the life of an individual, which in the case of mammalian animals can imply up to several decades (Deneris & Hobert, 2014). While yet relatively unexplored, seminal studies have proven the implication of distinct mechanisms in this process, including the self-regulation of terminal selectors induced in late developmental stages (Hobert, 2016), the *de novo* expression of cellular identity “keepers” (Aughey, 2023) and the involvement of distinct epigenetic factors maintaining appropriate epigenetic landscapes (Destain et al., 2024). For instance, alterations in histone post-translational modifications such as histone H3 tri-methylation in lysine 4 (Wang & Helin, 2025) and acetylation in lysine 27 (Tie et al., 2009), both associated with transcriptionally permissive states, or tri-methylation in lysine 9, a mark associated to transcriptionally repressed genes (Nicetto & Zaret, 2019), can compromise cell identity maintenance. This highlights the critical role that epigenetic effectors play in this process, nicely exemplified by the dramatic effects in acquired neuronal identity of the *Eed* gene elimination in dopaminergic and serotoninergic neurons (Toskas et al., 2022) or the *Ezh1/2* genes in striatal medium spiny neurons and Purkinje cells (von Schimmelmann et al., 2016), both part of the *Polycomb repressive complex 2* (PRC2) (Petracovici & Bonasio, 2021).

Supporting this idea, the genetic elimination of the lysine acetyltransferase type 3 (KAT3) proteins CBP and p300, two paralogues with a high level of functional redundancy as transcriptional co-activators (Lipinski et al., 2019), in excitatory neurons of the adult mouse brain severely compromised neuronal identity, causing a prominent neurological phenotype, electrophysiological alterations and a molecular signature associated to an undefined cellular identity state (Lipinski et al., 2020). This is in contrast with the more modest molecular and physiological consequences of the individual elimination of CBP or p300 in the same cells, highlighting the large functional redundancy between these paralogues (Lipinski et al., 2022). It is still unknown, however, whether these findings can be extrapolated to other CNS lineages and the possibility of reversing the phenotype by restating KAT3 activity.

The recently developed Proteolysis Targeting Chimera (PROTAC) dCBP-1 (Vannam et al., 2021) opens interesting possibilities for answering these questions. This heterobifunctional synthetic compound, whose design is based on the combination of the KAT3-selective bromodomain inhibitor GNE-781 (Romero et al., 2017) and a thalidomide-based cereblon (CRBN) E3-ligase ligand (Burslem & Crews, 2020), efficiently achieved a complete elimination of both KAT3 proteins within hours of treatment at nanomolar concentrations in different cancer cell lines, underscoring its suitability for combined CBP and p300 loss of function studies.

Here, we used dCBP-1 to characterize the molecular and physiological effects of acute KAT3 depletion in neuronal and glial cells with better temporal resolution than gene ablation approaches. Our results showed a rapid transcriptional dampening in both neuronal and glial primary cultures after acute KAT3 elimination, progressively affecting their lineage-specific cell-identity programs. These transcriptional effects early impacted physiological functions in neuronal and glial cultures, highlighting the constant dependance on KAT3 function of terminally differentiated CNS cell lineages. Interestingly, dCBP-1 washout was sufficient to restore physiological KAT3 levels, original transcriptome and several physiological functions in glial cells. These results open new avenues for the pharmacological manipulation of terminal neural cell identity, while raising concerns about the off-target effects that the therapeutical manipulation of KAT3 proteins can exert in the CNS.

## Results

### dCBP-1 mediated KAT3 elimination quickly impacts neuronal transcriptional identity

To interrogate the immediate effects of KAT3 depletion in neuronal cells we analyzed the effects of dCBP-1 in neuronal primary cultures from mouse embryonic hippocampal brain tissue. dCBP-1 treatment of mature cultures (day-in-vitro 14, DIV14) quickly depleted CBP and p300 protein levels, with a significant reduction already appreciable after only 3 hours of treatment (**Fig. 1a**). Similarly to the genetic ablation of KAT3 proteins in dKAT3ifKO mice primary cultures (Lipinski et al., 2020), dCBP-1 induced degradation of CBP and p300 in maturing neuronal cells produced a strong impairment of dendritic maturation, as observed by a reduced neurite length and number of branching points in dCBP-1 treated cells (**Supp. Fig. 1a-d**).

**Figure 1.**
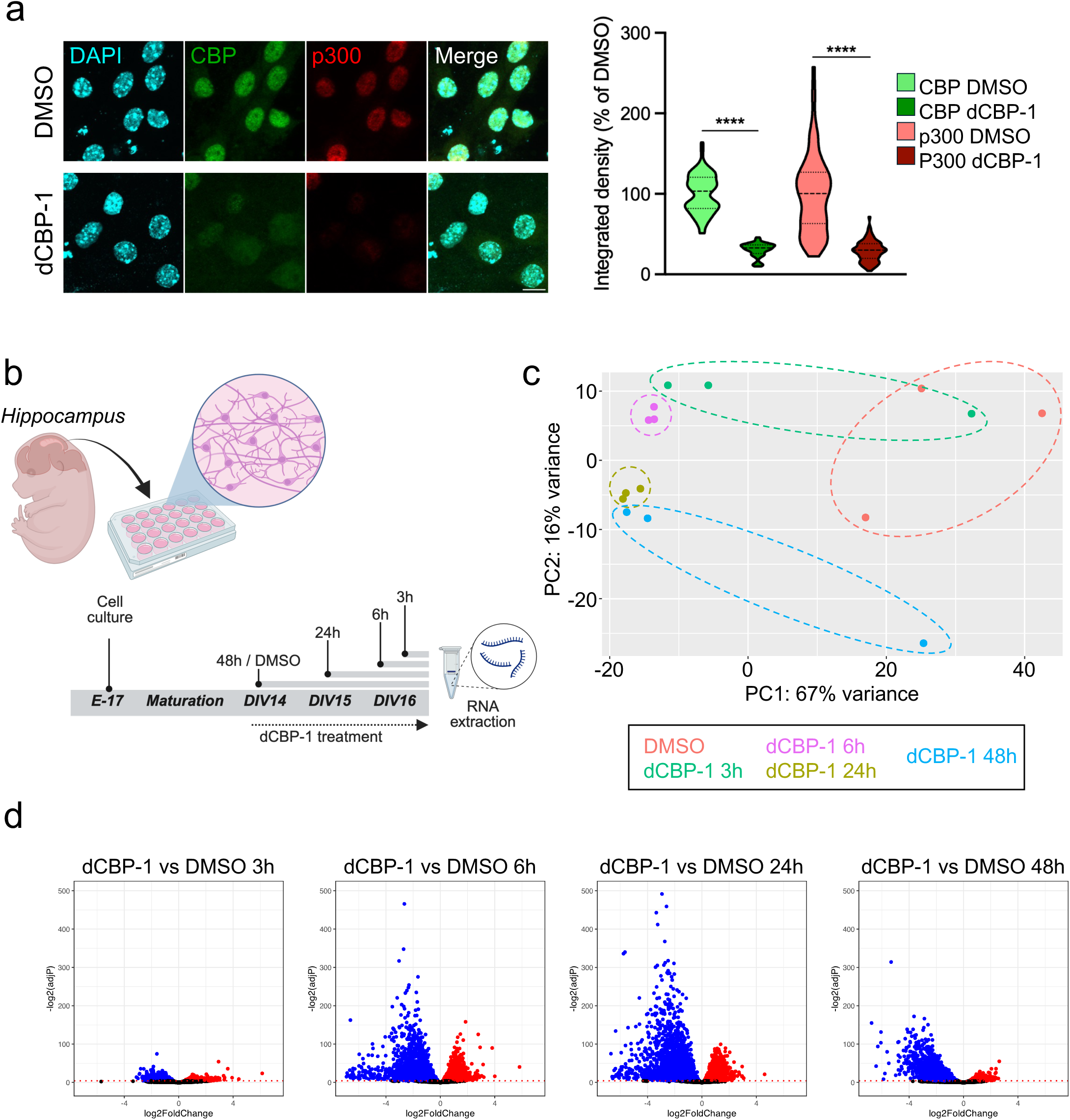
dCBP-1 transcriptional impact in neuronal primary cultures. **a.** On the left, representative confocal images of DAPI (cyan), CBP (green) and p300 (red) in neuronal cultures at DIV14 treated with dCBP-1 for 3 hours. Scale bar, 10 µm. On the right, violin plots showing CBP and p300 nuclear integrated density as % to DMSO controls in DMSO and dCBP-1 treated cells (N=4). ****P<0,0001 as compared with DMSO (Mann-Whitney test). **b.** Schematic representation of dCBP-1 time-course experiment using neuronal primary cultures from hippocampal embryonic mouse brain tissue. Created in Biorender. Fontana, T.(2024) . **c.** Principal Component Analysis (PCA) of neuronal dCBP-1 time-course RNA-seq data showing samples distribution (N=3) in PC1 and PC2 components using top 5000 genes capturing the largest source of variation. **d.** Volcano plots showing the log2FoldChange (x axis) and –log2(adjP) (y axis) of neuronal dCBP-1 RNA-seq time-course data from 3, 6, 24 and 48 hours of dCBP-1 treatment vs. DMSO. In blue and red, down- and up-regulated genes, respectively, between dCBP-1 and DMSO treated cells (adjP ≤ 0.05).

We decided then to evaluate the transcriptional dynamics arising from acute KAT3 elimination in mature neuronal cells by generating RNA-seq data in mature (DIV14) neuronal-enriched primary cultures after 3, 6, 24 and 48 hours of dCBP-1 treatment (**Fig. 1b**, **Supp. Fig. 1e** & **Supp. Table 1**). Principal Component Analysis (PCA) revealed a prominent effect of dCBP-1 in neuronal transcriptome from 3 hours in advance (**Fig. 1c**), with a predominant down-regulation of genes after 6 hours of treatment (**Fig. 1d**). CBP and p300 expression levels were not decreased by dCBP-1, but rather slightly increased at late time points probably as a compensatory response to the absence of KAT3 activity (**Supp. Fig. 1f**). Overall, we observed an increase in the percentage of reads at exonic regions and a reduction in intergenic ones, while no change was observed in the percentage of intronic reads (**Supp. Fig. 1g**) suggesting a more prominent affectation of distal regulatory regions which could fit with reduced activity at KAT3-regulated enhancer regions. In agreement with previous findings in dKAT3ifKO mice, down-regulated genes were particularly enriched in synaptic terms using SynGO when compared to up-regulated genes (**Supp. Fig. 2a,b**), including well-characterized neuronal genes such as *Neurod6* or *Syn1* (**Fig. 2a**). Supporting this, down-regulated genes significantly overlap with those identified by the inducible genetic elimination of KAT3 proteins in excitatory neurons, while little overlap was observed for up-regulated genes (**Fig. 2b**).

**Figure 2.**
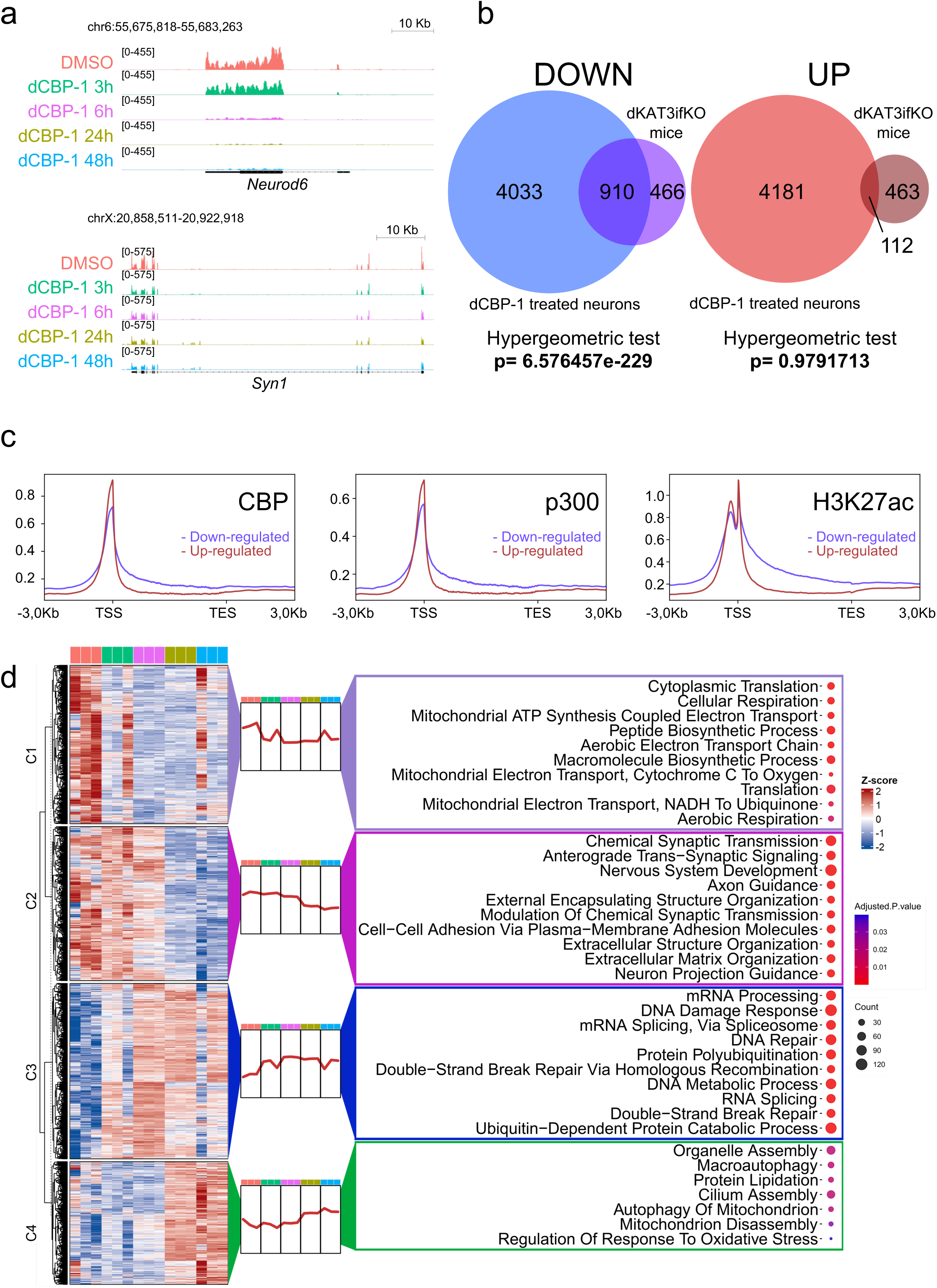
dCBP-1 progressively disrupts neuronal identity transcriptome in neuronal primary cultures. **a.** IGV genome tracks of dCBP-1 time-course RNA-seq data in neuronal primary cultures showing the progressive downregulation at the neuronal *Neurod6* and *Syn1* gene loci. **b.** Venn diagrams showing the overlap of genes down-(blue, left) and up-regulated (red, right) in dCBP-1 treated neuronal primary cultures and dKAT3ifKO RNA-seq data (Lipinski et al., 2020) from mouse hippocampus. Hypergeometric test shows the significance of the intersection with respect to chance probability. **c.** Gene body metaprofiles of mouse hippocampal CBP, p300 and H3K27ac ChIP-seq (Lipinski et al., 2020) signal enrichment in genes down-(blue) and up-regulated (red) in neuronal primary cultures treated with dCBP-1. **d.** On the left, K-means clustering heatmap showing normalized gene expression (z-score) of all differentially expressed genes identified in dCBP-1 time-course RNA-seq data using neuronal primary cultures. On the right, GO BP bubble plots showing enriched GO terms associated to the genes found in the different generated clusters. Bubble color indicates statistical significance (AdjPvalue) and size the gene counts associated to the term.

By analyzing the CBP and p300 chromatin binding and H3K27ac coverage in mouse hippocampal ChIP-seq data (Lipinski et al., 2020) at these two sets of genes, we observed that down-regulated genes presented a broader coverage of these marks across their gene body and preceding their TSS compared to up-regulated genes (**Fig. 2c**). This is line with previous reports suggesting a broader H3K27ac signal in super-enhancer regulated genes maintaining neuronal identity (Whyte et al., 2013).

To better understand the different transcriptional dynamics taking place upon dCBP-1 treatment, we applied k-means clustering to all differentially expressed genes, generating four clearly defined transcriptional patterns (**Fig. 2d**). Clusters 1 and 2 were enriched in downregulated genes, whereas clusters 3 and 4 contained mainly upregulated genes. Transcriptional changes followed a temporal progression, with cluster 1 exhibiting the earliest downregulation followed by cluster 2 and clusters 3 and 4 becoming upregulated from 3 and 24 hours, respectively. Functional enrichment analysis revealed that cluster 1 was associated with metabolism and cytoplasmic translation, cluster 2 with synaptic transmission and extracellular matrix organization, and clusters 3 and 4 with stress-response and proteasome-related pathways (**Fig. 2d**). SynGO analysis further showed that clusters 1 and 2 were linked to synaptic functions, specifically local protein synthesis and neurotransmission, respectively, whereas clusters 3 and 4 lacked synaptic enrichment (**Supp. Fig. 2c**). Notably, EP300 ChIP-seq enrichment was detected only in cluster 1 (Supp. Fig. 2d) using ChIP-X Enrichment Analysis (CheA), indicating that the earliest transcriptional changes may reflect direct KAT3-dependent regulation. These results evidence the constant need of KAT3 activity for maintaining the expression of neuronal transcriptional programs, probably due to their specific epigenetic landscape, highly demarcated by broad levels of H3K27ac.

### KAT3 elimination in neuronal primary cultures disrupts synaptic proteostasis and compromises synaptic function

The earliest transcriptional programs affected by KAT3 elimination in neuronal cells pointed to dysregulation of pathways involved in the synthesis and maintenance of synaptic proteins, suggesting that disruption of synaptic homeostasis may represent one of the earliest physiological consequences of KAT3 loss. To assess this hypothesis, we quantified the abundance of several synaptic proteins in mature primary hippocampal neurons treated with dCBP-1. Consistent with the relatively slow turnover of synaptic and dendritic proteins (Cohen et al., 2013), reductions in protein abundance were not immediately apparent but became detectable after 96 hours of KAT3 depletion (**Fig. 3a-c**). MAP2, an abundant structural dendritic protein with relatively slow turnover kinetics, exhibited a modest but significant reduction that became more pronounced with prolonged dCBP-1 treatment (**Fig. 3a**). Similarly, the presynaptic vesicular glutamate transporter vGlut1, and the postsynaptic scaffold protein Homer1 showed progressive decreases in abundance following KAT3 elimination (**Fig. 3b,c**). Both proteins are key components of mature excitatory synapses and are subject to regulation by neuronal activity and transcriptional programs (Yap & Greenberg, 2018). Other synaptic proteins, including Synapsin 1, exhibited a reduction in puncta number together with an apparent increase in protein aggregation or clustering (**Supp. Fig. 3a,b**). Together, these findings indicate that KAT3 elimination progressively disrupts the molecular composition of mature synapses. Rather than reflecting an acute depletion of individual proteins, the delayed and progressive loss of dendritic, presynaptic, and postsynaptic markers is consistent with impaired replenishment and maintenance of synaptic components, supporting a model in which KAT3-dependent transcription is continuously required to preserve synaptic proteostasis and structural integrity.

**Figure 3.**
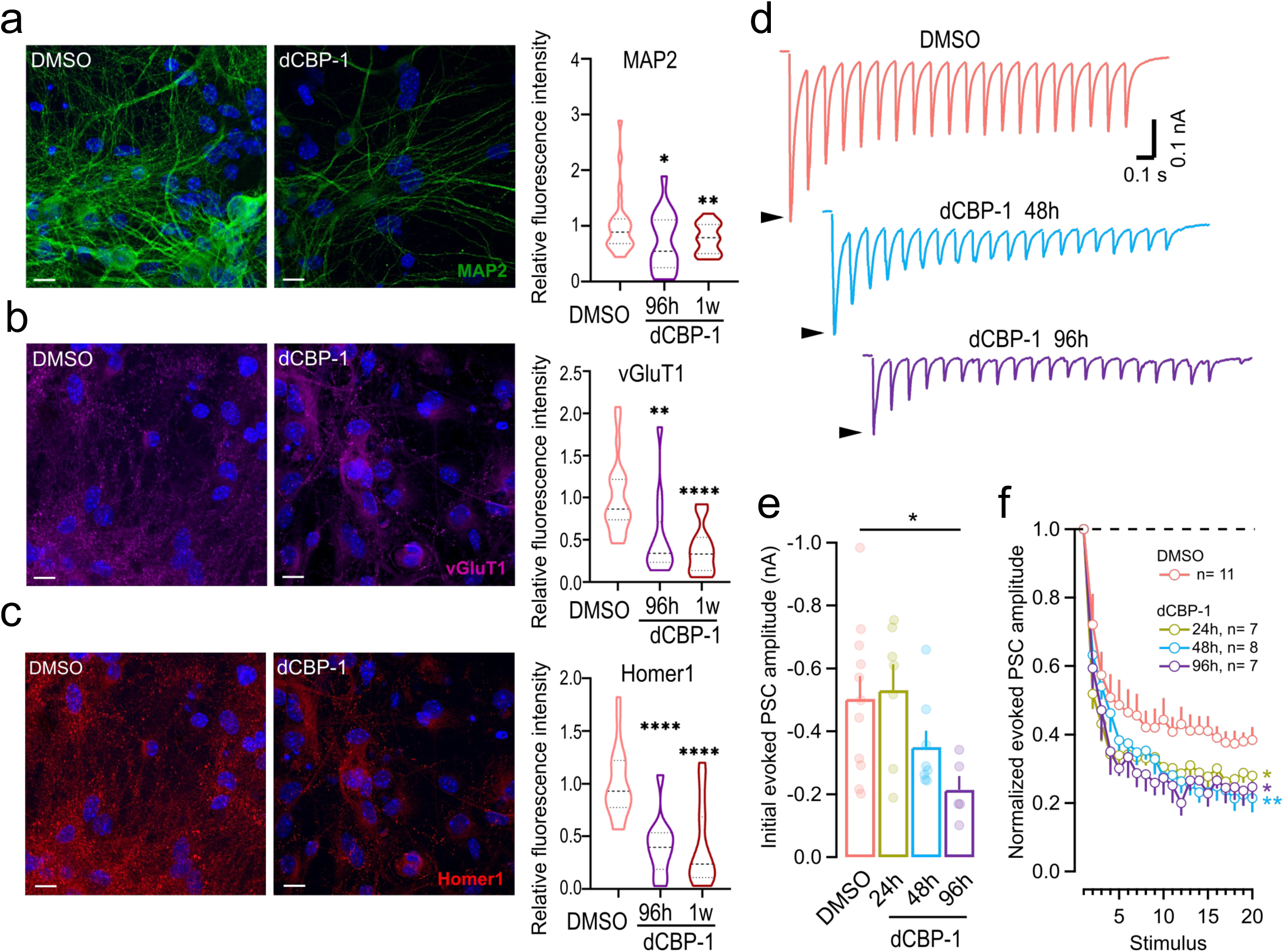
KAT3 elimination via dCBP-1 significantly impact synaptic composition and the electrophysiological properties of primary neuronal cultures. a-c. Representative confocal images of DIV14 hippocampal neurons treated with DMSO (control) or dCBP-1 for 96h or 1 week, followed by immunostaining for MAP2 (a), VGLUT1 (b), and Homer1 (c). Nuclei were counterstained with DAPI (blue). Scale bars, 10 μm. Right panels show quantification of fluorescence intensity normalized to DMSO-treated neurons (set to 1). Violin plots represent the distribution of values, with dashed lines indicating the median and quartiles. Quantifications were based on 27,17 and 16 images for MAP2; 23,12, and 17 images for vGLUT1; and 24,11, and 19 images for Homer1 in the DMSO, dCBP-1 96h, and dCBP-1 1-week groups, respectively, collected across 1–3 independent experiments. **d.** Average evoked postsynaptic current (ePSC) trains elicited by 10 Hz stimulation in control (DMSO, salmon) neurons and neurons treated with dCBP-1 for 48h (cyan) or 96h (violet). Black arrowheads indicate the amplitude of the first ePSC in each train. **e.** Mean amplitude of the first ePSC during 10 Hz stimulation, shown as bar graphs with individual recording values overlaid as scatter dot plots. **f.** Short-term plasticity assessed from ePSC amplitudes normalized to the initial ePSC during 10 Hz stimulus trains. Statistical significance in **a-c** was determined using two-tailed one-sample t-tests against a theoretical value of 1. In **e-f**, error bars indicate SEM. Statistical comparisons were performed using one-way ANOVA followed by Tukey test. *P < 0.05, **P < 0.01, ****P < 0.0001 versus DMSO.

To determine whether disruption of synaptic proteostasis was accompanied by functional deficits, we next examined action potential-evoked synaptic transmission in cultured hippocampal neurons following KAT3 elimination. Whole-cell recordings revealed a time-dependent reduction in synaptic strength, evidenced by a marked decrease in the amplitude of evoked postsynaptic currents (ePSCs) elicited by a single electrical stimulus applied to presynaptic neurons after 48h of dCBP-1 treatment (**Fig. 3d,e**). In contrast, neurons treated with dCBP-1 for 24 h exhibited synaptic strengths comparable to those of control neurons. However, assessment of short-term plasticity using 10-Hz stimulus trains revealed an earlier functional defect. After 24 h of dCBP-1 treatment, neurons displayed a pronounced reduction in the amplitude of successive ePSCs and stronger short-term depression compared with control cells, indicating impaired maintenance of neurotransmission during sustained neuronal activity. With longer treatment durations (48 and 96 h), both the initial synaptic response and subsequent responses during the stimulus trains were markedly reduced. Thus, defects in the ability to sustain synaptic transmission emerged before the substantial loss of synaptic structural proteins detected by immunofluorescence (**Fig. 3d,e**).

In parallel with the impairment of action potential-synchronous neurotransmission, dCBP-1-treated neurons exhibited a progressive increase in the frequency of spontaneous synaptic vesicle fusion events (**Supp. Fig. 3c,d**). This increase in spontaneous release, together with the reduced efficacy of evoked transmission, is consistent with a shift in the balance between activity-dependent and spontaneous neurotransmitter release and may reflect impaired maturation or maintenance of synaptic release mechanisms. Together, these electrophysiological findings indicate that KAT3 depletion rapidly compromises synaptic transmission, with functional abnormalities preceding the overt loss of synaptic structural components.

Collectively, these results demonstrate that KAT3 elimination initially impairs the functional capacity of mature synapses and is subsequently accompanied by progressive disruption of their molecular architecture. These findings highlight a critical role for KAT3-dependent transcriptional programs in maintaining synaptic proteostasis and preserving the structural and functional competence of mature excitatory synapses.

### dCBP-1 disrupts cell identity maintenance in glial cells

Previous results using dKAT3ifKO mice to genetically eliminate KAT3 proteins in glial primary cultures suggested that glial transcriptome is similarly sensitive to the lack of KAT3 activity (Lipinski et al., 2020), but genome-wide transcriptomic experiments providing confirmation of this were lacking at the time. To address that, we validated first that dCBP-1 was effectively depleting CBP and p300 in glial mouse primary cultures from 6 hours –the moment where we observed robust transcriptional changes in neurons– in advance (**Fig. 4a**). We generated then RNA-seq data at different dCBP-1 treatment time-points from glial primary cultures (**Fig. 4b** & **Supp. Fig. 4a**), which showed a progressive effect in glial transcriptome (**Fig. 4c**). Differential expression analysis unveiled a predominant gene down-regulation as observed in neuronal cultures (**Fig. 4d** & **Supp. Table 2**). As before, CBP and p300 expression levels remained relatively unchanged, with rather a slight increase in their expression (**Supp. Fig. 4b**). We also observed a subtle modification in the distribution of the reads across exons and introns, with the first ones gaining in mapped reads percentage in detriment of the seconds at late dCBP-1 treatment time-points (**Supp. Fig. 4c**). As in the case of neuronal cultures, down-regulated but not up-regulated genes in glial primary cultures presented a significant overlap with genes affected in dKAT3ifKO mouse hippocampal excitatory neurons lacking KAT3 proteins, suggesting that KAT3 proteins maintain the expression of common lineage determinant transcriptional programs in CNS lineages (**Fig. 4e**). As before, CBP, p300 and H3K27ac ChIP-seq data from adult mouse hippocampus showed a broader gene body enrichment of all three marks in down-regulated genes compared to up-regulated ones (**Supp. Fig. 4d**) – although more subtle than in previous neuronal analysis – suggesting that these genes might be also specially regulated by super-enhancer regulatory regions.

**Figure 4.**
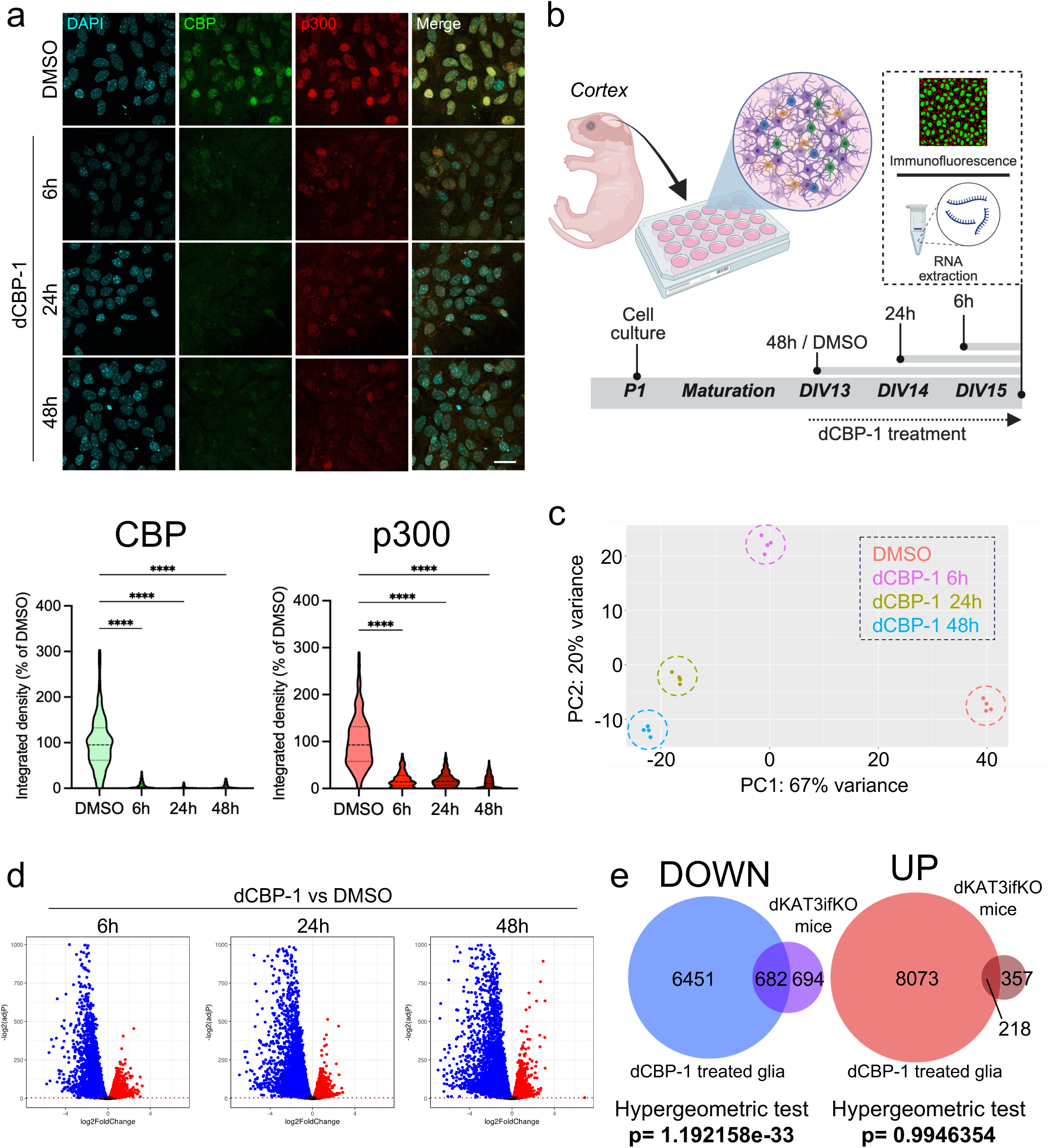
dCBP-1 transcriptional effects in glial mouse primary cultures. **a.** On the top, representative confocal images of DAPI (cyan), CBP (green) and p300 (red) in glial primary cultures treated with dCBP-1 for 6, 24 and 48 hours. Scale bar, 20 µm. On the bottom, violin plots showing CBP and p300 nuclear integrated density as % to DMSO controls in DMSO and dCBP-1 treated cells (N=4). ****P<0,0001 as compared with DMSO (Kruskal-Wallis test with Dunn’s multiple comparisons test). **b.** Schematic representation of dCBP-1 time-course experiment using glial primary cultures from post-natal (P1) cortical mouse brain tissue. Created in Biorender. Fontana, T.(2024). **c.** PCA of glial dCBP-1 RNA-seq time-course data showing samples distribution (N=4) in PC1 and PC2 components using top 5000 genes capturing the largest source of variation. **d.** Volcano plots showing the log2FoldChange (x axis) and –log2(adjP) (y axis) of glial dCBP-1 RNA-seq time-course data from 6, 24 and 48 hours of dCBP-1 treatment vs. DMSO. In blue and red, down- and up-regulated genes, respectively, between dCBP-1 and DMSO treated cells (adjP ≤ 0.05). **e.** Venn diagrams showing the overlap of genes down- (blue, left) and up-regulated (red, right) in dCBP-1 treated glial primary cultures and dKAT3ifKO RNA-seq data (Lipinski et al., 2020) from mouse hippocampus. Hypergeometric test shows the significance of the intersection with respect to chance probability.

When analyzing the GO enrichment in SynGo of down- and up-regulated genes, as observed in neurons, we observed that only down-regulated genes were enriched in processes related to synaptic functions, with a predominance in this case of terms related to Synapse Organization, while no pre- and post-synaptic terms were identified in contrast to neurons (**Supp. Fig. 5a,b** & **Supp. Fig. 2a,b**). To better dissect the different transcriptional dynamics taking place in these cells we applied k-means clustering to all differentially expressed genes and generated four different clusters with well-defined transcriptional profiles (**Fig. 5a**). Clusters 1 and 2 were enriched in downregulated genes, with cluster 1 showing a rapid and sustained decrease in expression and cluster 2 being affected mainly after 24 h. In contrast, clusters 3 and 4 contained upregulated genes, with cluster 3 exhibiting a transient early induction and cluster 4 increasing predominantly at later time points. GO analysis showed that downregulated clusters were enriched in glial-related functions, including cell migration, phagocytosis and Fc-gamma receptor signaling for cluster 1, and blood-brain barrier transport, cilium assembly and axon guidance for cluster 2 (**Fig. 5a**). Clusters 3 and 4, on the other hand, were primarily associated with stress response and metabolic pathways. For instance, some classic glial markers such as *Gfap* (astrocytes), *Olig1* (oligodendrocyte lineage) or *Cx3Cr1* (microglia) were identified within these clusters (**Supp. Fig. 5c**). ChEA analysis further revealed distinct regulatory signatures among downregulated genes. As observed in neurons, cluster 1 showed enrichment for EP300 binding sites, as well as SMRT- and IRF8-associated regulatory regions. By contrast, cluster 2 was enriched for SMARCA4 binding sites in oligodendrocytes, supporting a potential cooperative role between SMARCA4 and KAT3 proteins in controlling the expression of these genes (**Supp. Fig. 5d**).

**Figure 5.**
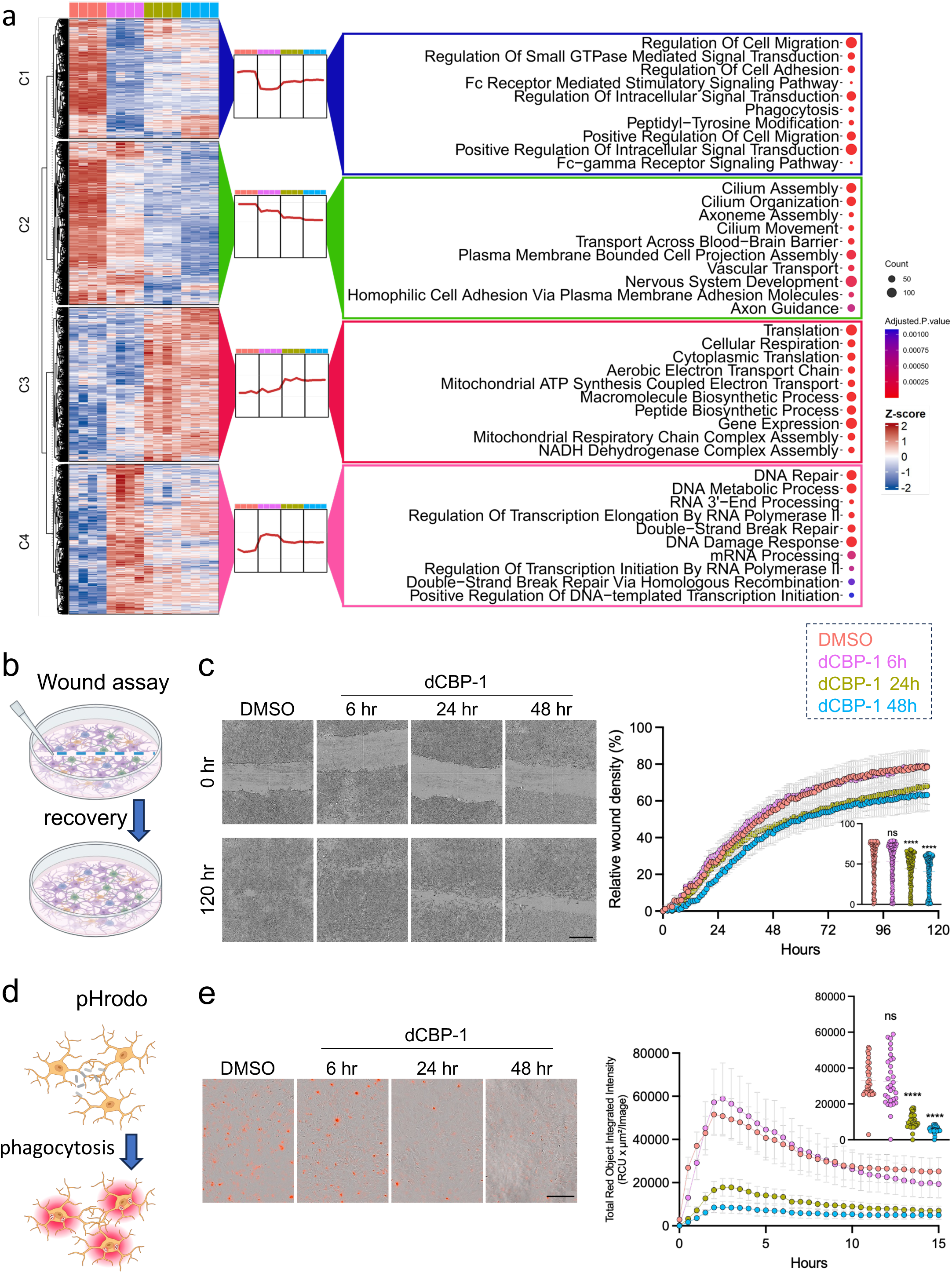
dCBP-1 abruptly impacts the glial identity transcriptome and physiological migration and phagocytic functions. **a.** On the left, K-means clustering heatmap showing normalized gene expression (z-score) of all differentially expressed genes identified in dCBP-1 time-course RNA-seq data using glial primary cultures. On the right, GO BP bubble plots showing enriched GO terms associated to the genes found in the different generated clusters. Bubble color indicates statistical significance (AdjPvalue) and size the gene counts associated to the term. **b.** Schematic representation of wound healing functional assay using glial primary cultures. Created in Biorender. Fontana, T.(2024). **c.** On the left, representative images of brightfield Incucyte® acquisitions of DMSO and dCBP-1 (6, 24 and 48 hours) treated cells at 0 and 120h post-wound. Scale bar, 500 µm. On the right, relative wound density quantification per hour or with all acquisition points together. Data is shown as mean ± SEM. ns, non-significant, ****P<0,0001 as compared with DMSO using Two-way Anova and Tukey’s multiple comparison test for mean group effect. **d.** Schematic representation of pHrodo fluorescence-based phagocytic functional assay using glial primary cultures. Created in Biorender. Fontana, T.(2024). **e.** On the left, representative images of brightfield and red pHrodo fluorescent signal from Incucyte® acquisitions of DMSO and dCBP-1 (6, 24 and 48 hours) treated cells at 15h after pHrodo addition to the culture. Scale bar, 200 µm. On the right, pHrodo signal quantification as Total Red Object Integrated Intensity (RCU x µm^2^ / Image) shown per hour or with all acquisition points together. Data is shown as mean ± SEM. ns, non-significant, ****P<0,0001 as compared with DMSO using Two-way Anova and Tukey’s multiple comparison test for mean group effect.

We wondered then if dCBP-1 treatment could affect some of the well-stablished glial physiological functions that appear to be affected at transcriptional level. As so, we first evaluated the cell migration capacity of dCBP-1 treated cells using a wound healing assay (**Fig. 5b**) (Liang et al., 2007) and live-imaging microscopy with Incucyte®. We observed a robust decrease in the migration capacity of glial cells treated with dCBP-1 from 24 hours in advance (**Fig. 5c**), suggesting that the deficient expression of transcriptional programs related to cell migration after KAT3 elimination quickly impacts glial migratory capacity. We also evaluated the phagocytic function of glial primary cultures treated with dCBP-1 using a pHrodo Bioparticles phagocytic assay (Takenouchi et al., 2009) (**Fig. 5d**). We observed a prominent reduction in pHrodo integrated intensity from 6 hour of dCBP-1 treatment in advance (**Fig. 5e**) resulting from a reduced number of cells with phagocytic capacity (**Supp. Fig. 5e**) and a reduction of the average intensity per cell (**Supp. Fig. 5f**). This suggests that the transcriptional shutdown of phagocytosis and cell migration related gene programs upon KAT3 elimination has a hasty effect in these physiological glial functions. Altogether, these results highlight the crucial role of KAT3 proteins as cell identity keepers in the glial lineage, being crucial for maintaining the functionality of specialized glial physiological processes.

### KAT3 elimination leads to a progressively convergent transcriptional state in CNS neural lineages

Our results indicated that KAT3 degradation in neuronal and glial lineages progressively leads to the downregulation of transcriptional programs associated with their specific acquired cell identities. This made us wonder if, while reducing those cell-specific transcriptional programs the similarity between these lineages might arise, progressively leading them to a cell identity state where cells barely express specific cell markers differentiating one from each other. To assess this possibility, we integrated the two time-course RNA-seq series and analyzed the progressive alterations occurring in their specific cell identity programs. We took advantage of previous cell identity markers identified using hippocampal single-nuclei RNA-seq data from adult mouse brain to address this question (Lipinski et al., 2020). We validated first that in basal conditions each primary culture significantly expressed in a higher extent the markers associated to their specific lineage (**Fig. 6a**). For instance, *Neurod6* and *Mbp*, two well characterized genes prominently expressed in neuronal and glial cells, respectively, were highly enriched in their expected primary culture, but not in the other (**Fig. 6b**). PCA analysis of integrated RNA-seq data revealed a common trajectory in the transcriptional effects caused by dCBP-1 on both lineages, suggesting that KAT3 elimination renders them to a similar transcriptional state when losing their original identity (**Fig. 6c**). By analyzing the expression of hippocampal cell-specific markers in both time-course datasets, we observed a drastic progressive depletion of their cell identity markers (**Fig. 6d**), reaching a similar indefinite cell identity state in both cellular lineages (**Supp. Fig. 6a**).

**Figure 6.**
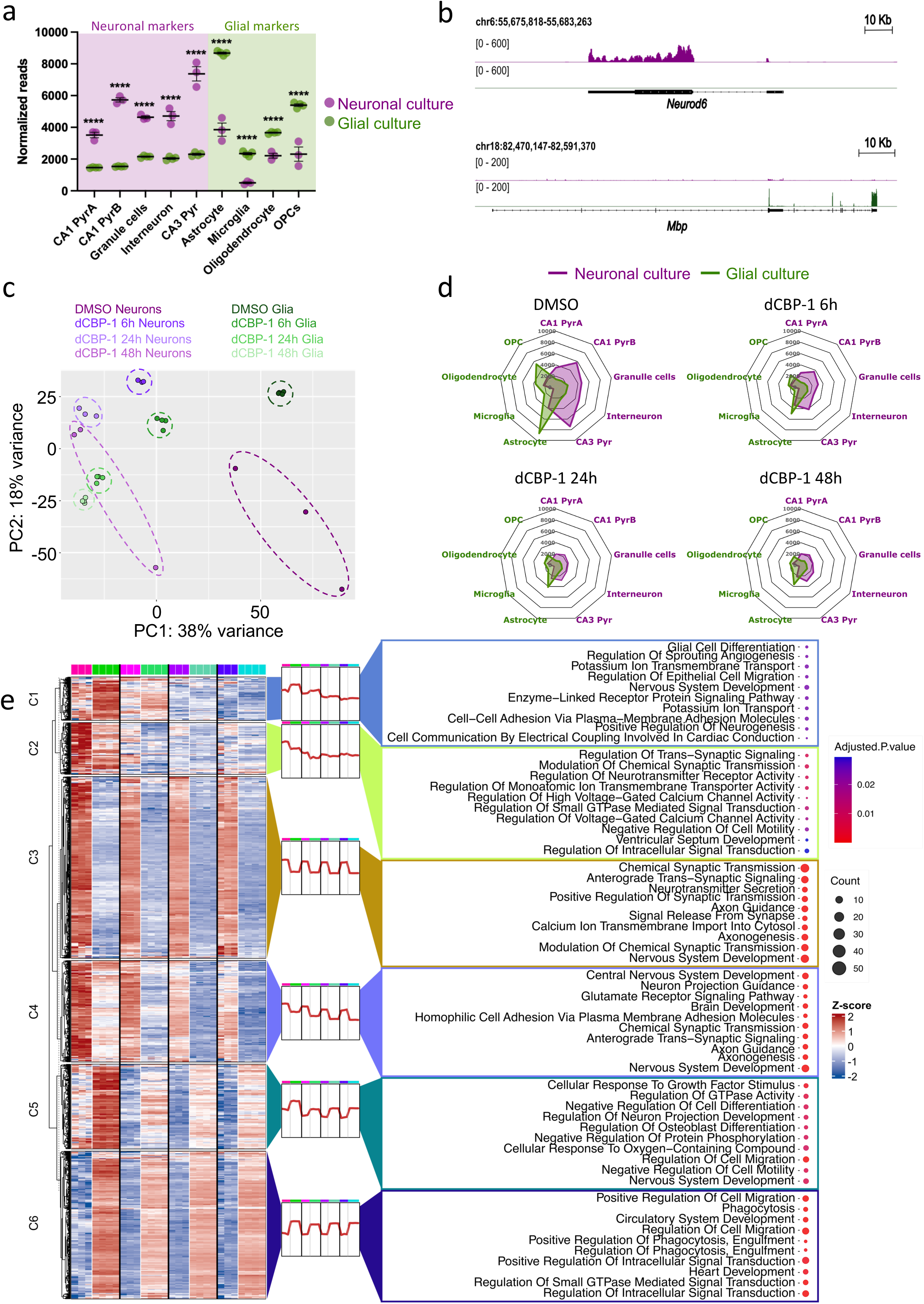
dCBP-1 treatment progressively leads to a cell-identity indefinite state. **a.** Hippocampal cell identity markers normalized mean expression in integrated RNA-seq data from neuronal (purple) and glial (green) mouse brain primary cultures. ****P<0,0001 comparing neuronal versus glial cultures expression using Two-way Anova and Bonferroni’s multiple comparison test. **b.** IGV genome tracks of DMSO treated controls in neuronal (purple) and glial (green) integrated RNA-seq data from dCBP-1 time-course showing the neuronal and glial enrichment of *Neurod6* and *Mbp* gene expression, respectively. **c.** PCA of integrated neuronal (purple tonalities, N=3) and glial (green tonalities, N=4) RNA-seq time-course data showing samples distribution in PC1 and PC2 components using top 5000 genes capturing the largest source of variation. **d.** Radar plots showing the mean expression of neuronal (purple fonts) and glial (green fonts) markers from mouse hippocampal brain tissue (Lipinski et al., 2020) in neuronal (purple) and glial (green) RNA-seq time-course data of DMSO or dCBP-1 (6, 24 and 48 hours) treated cells. **e.** On the left, K-means clustering heatmap showing normalized gene expression (z-score) of all cell identity markers from mouse hippocampal brain tissue using neuronal and glial integrated dCBP-1 time-course RNA-seq data. On the right, GO BP bubble plots showing enriched GO terms associated to the genes found in the different generated clusters. Bubble color indicates statistical significance (AdjPvalue) and size the gene counts associated to the term.

We wondered if these markers were affected following different temporal dynamics depending on their specific function or enrichment in CBP and p300. To answer this, we applied k-means clustering to all hippocampal cell markers previously analyzed using our two sets of RNA-seq data (**Fig. 6e**). We generated six distinct clusters attending to their enrichment in one cell type or the other and to their different temporal transcriptional dysregulation upon dCBP-1 treatment. Clusters 2, 3 and 4 were more expressed in neuronal cultures while clusters 1, 5 and 6 presented a higher expression in glial cells. Clusters 1 and 2 were highly enriched in glial and neuronal cells, respectively, and were early impacted by dCBP-1 treatment, reaching at 48 hours a similar expression between both cell types. On the other hand, clusters 4 and 5 presented a similar level of lineage specific expression at basal conditions but displayed a smaller reduction in their expression throughout the time-course. Finally, clusters 3 and 6 were also prominently enriched in neuronal and glial cultures, respectively, in DMSO treated cells, but their expression was only slightly reduced in response to dCBP-1, even at late time-points. GO BP analysis confirmed the specificity of these gene programs, showing classically associated functions of glial and neuronal cells in the different clusters according to their enrichment in one lineage or the other (**Fig. 6e**).

Analyzing the enrichment of CBP, p300 and H3K27ac in hippocampal ChIP-seq data we observed that clusters 2 for neurons and cluster 1 for glia, which were the earlier impacted by the treatment, presented a higher coverage of these three marks along their gene bodies (**Supp. Fig. 6c**), suggesting that their earlier sensitivity to KAT3 elimination might be related to their specific epigenetic landscape. This loss of cell identity markers occurred without the re-induction of developmental transcriptional programs, as depicted by the general decreased expression of mouse cortical developmental modules recently generated using longitudinal single-cell RNA-seq experiments from mouse developmental and early post-natal cortical tissue (Gao et al., 2025) (**Supp. Fig. 6d-e**). Overall, our data suggests that in the absence of KAT3 proteins different CNS lineages tend to converge to a similar cell identity state characterized by the depletion of cell-specific markers following distinct temporal dynamics according to their initial epigenetic landscape.

### KAT3 re-statement after cell identity disruption reestablishes glial transcriptome and physiological functions

Taking advantage of our pharmacological approach and the recurrent need for media exchange of glial cultures, we decided to conduct a KAT3 recovery experiment (**Fig. 7a**) to evaluate the possible reversibility of this process. We validated first that, after a single administration of dCBP-1, CBP and p300 protein levels were completely depleted for at least 72 hours in our glial cultures (**Supp. Fig. 7a-b**), and we set up that interval for every media exchange in the recovery experiment. We validated that with this configuration, after one week of dCBP-1 treatment, KAT3 proteins were fully eliminated in glial primary cultures (**Supp. Fig. 7c**). In parallel cultures, we conducted an extra week of dCBP-1 treatment (dCBP-1 2w) or DMSO in control (DMSO 2w) or previously dCBP-1 treated (Recovery) cells with every three-day media exchange. Inmunofluorescence analyses evidenced that CBP and p300 protein levels could be restored to basal levels upon dCBP-1 washout (**Fig. 7b**) with this configuration.

**Figure 7.**
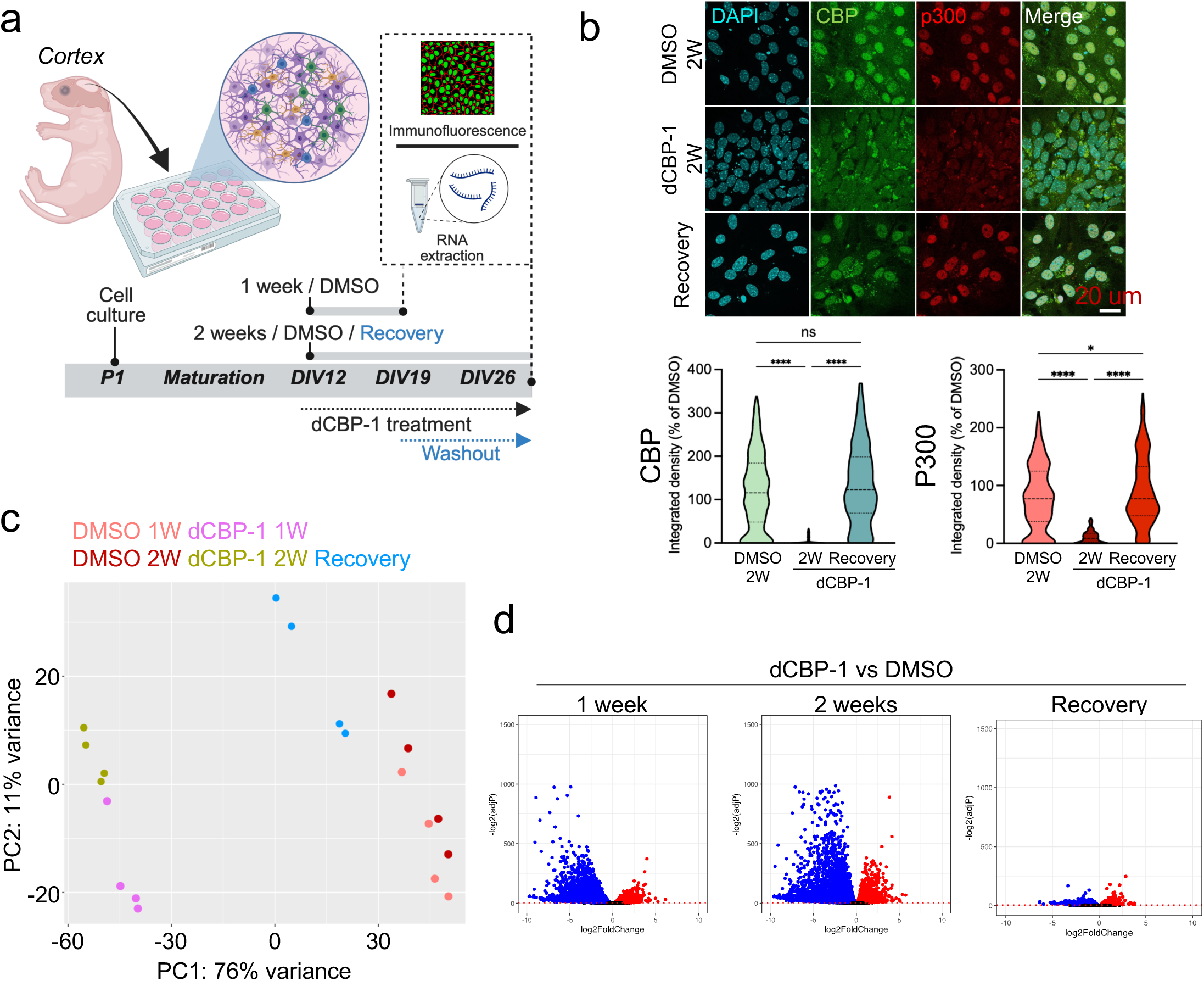
dCBP-1 washout restores KAT3 levels and glial transcriptome. **a.** Schematic representation of dCBP-1 recovery experiment using glial primary cultures from post-natal (P1) cortical mouse brain tissue. Created in Biorender. Fontana, T.(2024). **b.** On the top, representative confocal images of DAPI (cyan), CBP (green) and p300 (red) in glial primary cultures treated for 2 weeks with DMSO (DMSO 2W), dCBP-1 (dCBP-1 2W), or 1 week with dCBP-1 and the last week with DMSO (Recovery). Scale bar, 20 µm. On the bottom, violin plots showing CBP and p300 nuclear integrated density as % to DMSO controls (DMSO 2W) in DMSO 2W, dCBP-1 2W and Recovery conditions (N=4). ns, non-significant, ****P<0,0001 as compared with DMSO 2W (Mann-Whitney test). **c.** PCA of glial dCBP-1 recovery RNA-seq data showing samples distribution (N=4) in PC1 and PC2 components using top 5000 genes capturing the largest source of variation. **d.** Volcano plots showing the log2FoldChange (x axis) and – log2(adjP) (y axis) of glial dCBP-1 recovery RNA-seq data from 1 week, 2 weeks and Recovery conditions vs. DMSO. In blue and red, down- and up-regulated genes, respectively, between dCBP-1 and DMSO treated cells (adjP ≤ 0.05).

We then evaluated the impact in glial transcriptome of this transient KAT3 elimination by generating RNA-seq data in these different experimental conditions (**Fig. 7c**, **Supp. Fig. 7d** & **Supp. Table 3**). PCA analysis showed a clear separation of DMSO and dCBP-1 treated cells, with PC1 explaining 76% of the variability between these samples. Remarkably, Recovery samples clustered with DMSO treated cells, although did not fully overlap (**Fig. 7c**). This was also clearly evidenced by the global Spearman correlation coefficient between samples, which clustered together DMSO and Recovery conditions in contrast with dCBP-1 treated samples (**Supp. Fig. 7e**). Differential expression analysis revealed a prominent transcriptional dysregulation in response to dCBP-1 in both, 1-week and 2-week treated samples, and almost a complete reversion of these changes in the Recovery condition (**Fig. 7d**). These changes, in contrast with previous findings at short time points, didn’t affect the global distribution of reads among exons, introns or intergenic regions (**Supp. Fig. 7f**).

To better dissect the transcriptional dynamics leading to this recovery, we generated three different lists of genes according to their recovery state (**Fig. 8a** and **Sup. Fig. 8a**): fully recovered genes, partially recovered genes and not recovered genes (see Methods section for details). GO BP analysis of these different sets of genes showed how fully and partially recovered genes were enriched in glial processes previously identified to be affected by dCBP-1 treatment including ‘Cell migration’, ‘Transport Across Blood-Brain Barrier’ or ‘Phagocytosis’ (**Fig. 8b**).

**Figure 8.**
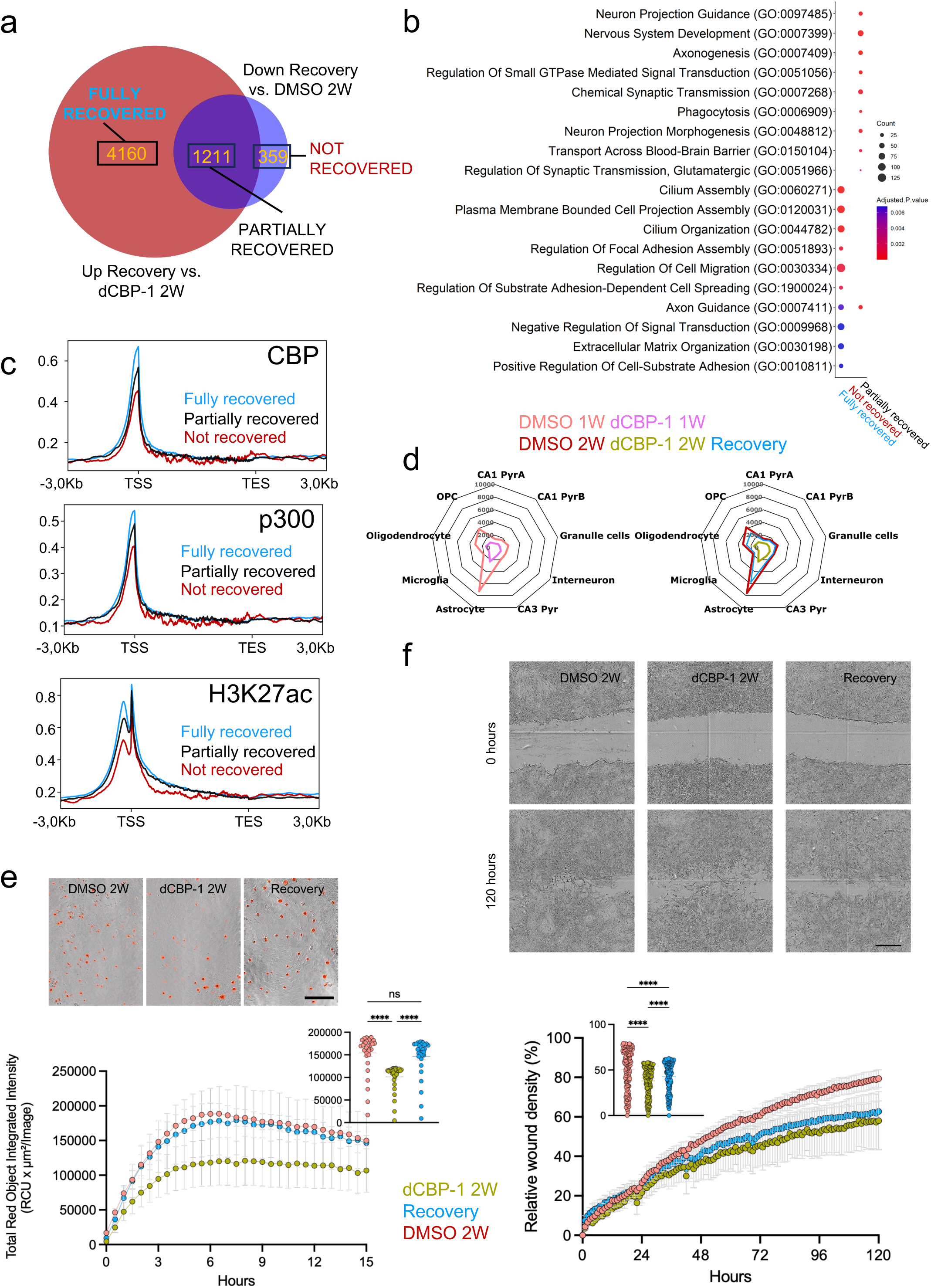
dCBP-1 washout restores glial identity transcriptome and partially restores physiological functions. **a.** Venn diagram Venn showing the overlap of genes significantly up-regulated between Recovery and dCBP-1 2W (red) and genes significantly down-regulated between Recovery and DMSO 2W (blue) generating three different categories of genes, fully recovered, partially recovered, and not recovered genes. **b.** GO BP bubble plots showing enriched GO terms associated to the genes belonging to the fully recovered, partially recovered and not recovered transcriptional categories. Bubble color indicates statistical significance (AdjPvalue) and size the gene counts associated to the term. **c.** Gene body metaprofiles of mouse hippocampal CBP, p300 and H3K27ac ChIP-seq (Lipinski et al., 2020) signal enrichment in fully recovered (blue), partially recovered (black) and not recovered (red) gene lists from dCBP-1 recovery RNA-seq experiment. **d.** Radar plots showing the mean expression of glial markers from mouse hippocampal brain tissue (Lipinski et al., 2020) in 1-week DMSO and dCBP-1 (left) and 2-weeks DMSO, dCBP-1 and Recovery (right) RNA-seq data. **e.** On top, representative images of brightfield Incucyte® acquisitions of DMSO 2W and dCBP-1 (2W and Recovery) treated cells at 0 and 120h post-wound. Scale bar, 500 µm. On the bottom, relative wound density quantification per hour or with all acquisition points together. Data is shown as mean ± SEM. ****P<0,0001 as compared with DMSO using Two-way Anova and Tukey’s multiple comparison test for mean group effect. **f.** On top, representative images of brightfield and red pHrodo fluorescent signal from Incucyte® acquisitions of DMSO 2W and dCBP-1 (2W and Recovery) treated cells at 15h after pHrodo addition to the culture. Scale bar, 200 µm. On the bottom, pHrodo signal quantification as Total Red Object Integrated Intensity (RCU x µm^2^ / Image) shown per hour or with all acquisition points together. Data is shown as mean ± SEM. ns, non-significant, ****P<0,0001 as compared with DMSO using Two-way Anova and Tukey’s multiple comparison test for mean group effect.

Aiming to elucidate the molecular basis leading to these distinct recovery dynamics we analyzed different possibilities. We first noticed that transcriptional dampening in Partially recovered genes was greater than in Fully recovered ones, suggesting that maybe Partially recovered genes needed more time to fully achieved DMSO control levels (**Supp. Fig. 8a**). We also considered the possibility that cells could re-acquire their original identity transcriptome following their initial developmental programs, but we didn’t observe a particular enrichment for cortical developmental modules of the different gene sets that pointed on that direction (**Supp. Fig. 8b**). We then wondered if their original epigenetic landscape was better correlating with their identity re-acquisition in response to KAT3 restatement. To answer this, we analyzed the gene body enrichment of CBP, p300 and H3K27ac in the three different sets of genes (**Fig. 8c**). The level of enrichment for the three marks followed the same pattern, with a higher enrichment in fully recovered genes, followed by partially recovered genes and not recovered ones, suggesting that their original epigenetic landscape might be a critical factor explaining these differences. ChEA analysis reinforced this idea, as fully recovered genes were the only ones showing an enrichment in EP300 binding sites identified in the mouse forebrain (**Supp. Fig. 8c**). Finally, we also consider the possibility that different glial lineages might recover at distinct paces, so we analyzed the expression of the previously used glial identity markers in the three sets of genes (**Supp. Fig. 8d**). While OPC identity genes were relatively even between the different groups, we observed that astrocyte and oligodendrocyte markers were especially enriched in fully and partially recovered categories, while microglial markers accumulated more in not recovered ones, suggesting that the differences in recovery dynamics might also be cell-type dependent. Despite this, we observed a general identity rescue in the Recovery condition when analyzing hippocampal identity markers as compared to dCBP-1 treated cells (**Fig. 8d**), including the expression of *Gfap*, *Olig1* and partially *Cx3cr1* (**Supp. Fig. 8**).

We wondered if this transcriptional recovery could be enough to functionally restore the cell migration and phagocytic functions of glial cells. Phagocytic function assessed by pHrodo fluorescence intensity in response to its internalization appeared to be fully normalized in Recovery cells (**Fig. 8e**). This recovery was even more striking considering that the number of phagocytic cells was still significantly depleted in the Recovery condition, and it was rather originated by even an exacerbated average phagocytic capacity of individual cells compared to DMSO or dCBP-1 treated ones (**Supp. Fig. 8f,g**). Wound healing assay also showed a slight but significant improvement in the wound healing capability in the Recovery condition compared to dCBP-1 treated ones, although still far from reaching control levels (**Fig. 8e**). Altogether, these results evidenced that cell identity maintenance in the glial lineage can be dynamically tuned by modulating KAT3 protein levels, affecting both molecular and physiological processes, providing an interesting molecular switch with potential biomedical applications.

## DISCUSSION

Understanding the underlying mechanisms governing cellular specification during mammalian brain development has been one of the major challenges of neurobiology in the last decades (Gao et al., 2025; Lodato & Arlotta, 2015). With the disentanglement of the molecular basis driving this orchestrated cell diversification, it became unquestionable that terminal selectors play a crucial role in this process by governing the expression of terminal effector gene programs (Hobert & Kratsios, 2019). However, it was not until pivotal studies proved that terminal selectors are not only crucial for neuronal identity acquisition but also at post-developmental stages that the fundamental role of cell identity maintenance in the CNS was not started to be foresaw (Doitsidou et al., 2013; Flames & Hobert, 2009; Laguna et al., 2015). This opened fundamental questions in the field, as it blurred the frontier between developing and differentiated cells, questioning the concept of identity perse (Morris, 2019). Can we attribute an identity to a cell if this one is in constant evolution or what we call identity is nothing else than a lesser dynamic state of what a cellular precursor was in earlier developmental stages? The study of these fundamental questions brought to light additional molecular mechanisms contributing to this active identity maintenance process, including the induction of different transcription factors dedicated to this purpose or the action of distinct epigenetic effectors (Deneris & Hobert, 2014).

Based on previous evidence involving KAT3 proteins in the cell identity maintenance of excitatory neurons of the adult mouse brain (Lipinski et al., 2020), our study aimed to interrogate the dynamic molecular changes immediately arising after a major identity maintenance insult in the two main lineages composing the CNS, neurons and glial cells. With our time-course RNA-seq data using the dCBP-1 KAT3 targeting PROTAC we have shown the susceptibility and resilience of different transcriptional cell identity programs specific to both lineages. These results, together with our functional data provide a precise temporal picture of the molecular and physiological consequences of perturbing cell identity maintenance in the CNS. For instance, our time-course synaptic immunostainings and electrophysiological recordings in dCBP-1 treated neuronal cells clearly demonstrated that KAT3 activity is essential for the proper proteostatic balance of synaptic components, but not crucial for evoked excitatory synaptic transmission, which could be still observed days after KAT3 elimination.

At the molecular level, and in line with previous evidence pinpointing the enrichment of CBP and p300 in H3K27ac wide genomic domains (Consortium, 2013; Ramos et al., 2010), our data suggest that gene programs primarily impacted by KAT3 depletion fit well with the classical description of super-enhancer regulated genes (Whyte et al., 2013). In this regard, we observed a prominent difference in the genomic distribution of our RNA-seq reads between neuronal and glial data, with the first presenting a higher read coverage at intergenic regions. This is in line with the more complex epigenomic landscape classically associated to neuronal cells (Closser et al., 2022; Zylka et al., 2015), that could contribute to the retrieval of a higher number of enhancer RNAs frequently located in non-coding intergenic regions (Kim et al., 2010). Interestingly, we observed a reduction of intergenic reads in favor of exonic ones in our neuronal RNA-seq data upon dCBP-1 treatment, suggesting that KAT3 depletion could compromise the proper expression of these enhancer RNAs specifically in the neuronal lineage.

A major finding of our study is the fact that this lost identity could be re-stated both at transcriptional and functional level by recovering KAT3 activity, suggesting that complementary epigenetic mechanisms might remain in place for the re-activation of KAT3-dependent cell identity transcriptional programs. This transcriptional recovery didn’t appear to follow lineage-specific developmental trajectories but rather seemed to respond to the original epigenetic landscape and the extent of transcriptional dampening of affected genes. It is tempting to speculate that cell reprogramming strategies might benefit from cells being in this transient “cell identity limbo”, improving similar previous attempts using KAT3 inhibitors (Ebrahimi et al., 2019).

Our results, while primarily contributing to our understanding of fundamental biological principles governing cell identity maintenance in the CNS can also be of relevance for better understanding the molecular basis of neurodevelopmental and neurodegenerative disorders. Despite no clinical case has yet been reported in which both KAT3 paralogues’ expression is compromised - and it is not prone to happen due to neural tube closure defects observed in homozygotic KAT3 mutation carriers (Bhattacherjee et al., 2009; T. P. Yao et al., 1998) -, heterozygotic mutations in *CREBBP* or *EP300* cause Rubinstein Taybi Syndrome (RSTS), a rare neurodevelopmental intellectual disability disorder characterized by distinctive facial features, microcephaly and cognitive impairment (Tanaka et al., 1997; Van Gils et al., 2021). Considering that at particular developmental stages certain tissues might only express one of the two paralogues (Chen et al., 2009), similar scenarios to our experimental setup in which KAT3 activity is temporary compromised might arise at specific developmental stages and cell types in RSTS individuals. However, a more similar scenario to our experimental setup could be found in neurodegenerative disorders in which KAT3 protein activity is compromised at adult stages due to disease-associated pathological traits. In this regard, Huntington’s disease (HD), a neurodegenerative disease caused by the aberrant expansion of a CAG triplet within the exon 1 of the *HTT* gene (Donaldson et al., 2026), could be one of the more resemblant situations, as CBP (and probably p300) is sequestered within mutant huntingtin aggregates, greatly impacting KAT3 activity (Matsumoto et al., 2006). Supporting this, transcriptional and epigenetic perturbations of neuronal cell identity genes have been extensively described in HD (Alcalá-Vida et al., 2021; Langfelder et al., 2016). Interestingly, mutant huntingtin cell-specific lowering using an AAV encoded Zinc Finger Protein (ZFP) targeting huntingtin CAGs resulted in the disassembly of mutant huntingtin aggregates, probably freeing sequestered CBP, and was able to restore neuronal identity transcriptome (Gangwani et al., 2023), nicely supporting our findings regarding the reversibility of the process.

Importantly, our data also serve as a cautionary message for KAT3 targeting strategies in the cancer field (Nicosia et al., 2023; Wu et al., 2025), underscoring the huge potential off-target effects of these approximations in cellular lineages where identity maintenance strongly relies on KAT3 activity such us in the CNS.

Overall, our study provides a great insight about the molecular underpinning of KAT3-dependent cell identity maintenance in the CNS, providing for the first time a pharmacological switch to easily modulate this process with great potential biomedical and fundamental biology applications.

## Acknowledgements

We thank the personnel of the Mouse facility, and the Omics and microscopy core services at the Instituto de Neurociencias for their assistance. The authors would like to acknowledge the Genomics Unit at the CRG for assistance with library preparation and sequencing. We also thank R. Olivares and C. Racovac for technical assistance. J.Z.L. was recipient of a fellowship from the CIACIF Generalitat Valenciana Program. S.N. was recipient of a fellowship from the Spanish Ministry of Science and Innovation (MICINN). V.E.A.M is funded by a SEJIGENT (CISEJI/2023/40) Project from the Generalitat Valenciana. R.A.V was recipient of a ‘la Caixa’ Junior Leader Postdoctoral Fellowship (LCF/BQ/PI22/11910006) and a Ramón y Cajal contract (RYC2024-048542-I) from the Agencia Estatal de Investigación. The work of F.J.L-M and T.L.-H was supported by the Agencia Estatal de Investigación (PID2022-141685NA-I00, PID2025-168433NB-I00, F.J.L.-M.; PID2021-128845NA-I00, T.L.-H.), by the Max Planck Partner Group Program (F.J.L.-M.) and by the Fundación Ramón Areces (T.L.-H.) A.B. research is supported by grants PID2023-148442NB-I00 by MICIU/AEI/10.13039/501100011033 co-financed by ERDF, FCAIXA HR22-00394 from Fundación LaCaixa, and CIPROM/2023/15 from the Generalitat Valenciana. R.A.V research is supported by a Fundación LaCaixa Junior Leader Postdoctoral Fellowship LCF/BQ/PI22/11910006 and a SEJIGENT Project from the Generalitat Valenciana CISEJI/2023/40. The Instituto de Neurociencias is a “Centre of Excellence Severo Ochoa” (CEX2021-001165-S).

## Author contributions

Conceptualization, J.Z.L, A.B. and R.A.V; Methodology, J.Z.L, V.E.A.M, F.J.L.M. and T.L.H; Software, J.Z.L and S.N.; Experimental work, J.Z.L, V.E.A.M, F.J.L.M. and T.L.H; Data Curation and Visualization, J.Z.L, S.N, V.E.A.M, F.J.L.M, T.L.H and R.A.V.; Writing - Original Draft, J.Z.L, V.E.A.M and R.A.V; Data discussion and Manuscript Revision A.B.; Funding Acquisition, R.A.V.

## Declaration of generative AI and AI-assisted technologies in the writing process

The authors used Microsoft Copilot (Microsoft Corporation) to assist with language editing, text refinement, and identification of relevant scientific literature. All retrieved references were independently verified by the authors, who take full responsibility for the final content of the manuscript.

## Methods

### Mice

Animals were maintained according to the Spanish and European regulations, and the followed experimentation were approved by the Animal Welfare Committee at the Instituto de Neurociencias and the CSIC Ethical Committee. For primary cultures ICR embryos and P0-P1 mice were used to generate neuronal and glial primary cultures, respectively.

### Neuronal primary cultures

Primary hippocampal neurons were obtained from E17 mouse embryos. Briefly, hippocampi were micro-dissected, dissociated in Trypsin 0.05% and seeded in poly-D-lysine pre-treated p24 well plates with or without a 13 mm glass coverslip depending on downstream experiments at 110.000 neurons/well and maintained in Neurobasal medium supplemented with 1x B-27 (Gibco), Glutamine (10mM, Gibco) and Pen/Strep (500U/ml, Gibco) as previously described in (Benito et al., 2011).

### Glial-enriched primary cultures and dCBP-1 administration

Primary cortical glial cultures were prepared from postnatal P0-P1 cortices. Briefly, mouse cortices were micro-dissected and mechanically homogenized with successively thinner glass Pasteur pipettes until obtaining a homogeneous solution. Cellular suspensions were seeded in poly-D-lysine pre-treated p24 well plates with 13 mm glass coverslip or directly on plastic depending on downstream experiments and maintained in DMEM with pyruvate (Merck) supplemented with 10% FBS (Gibco) and Pen/Strep (500U/ml, Gibco) until reaching cellular confluence (DIV12-13) as previously described (Lipinski et al., 2020) .

### dCBP-1 treatment

dCBP-1 PROTAC (Merck) was dissolved in DMSO to generate a stock at 5mM concentration and diluted to 500nM when applied to neuronal and glial primary cultures generating a 0.25 nM working solution for using 1 μl volume per 500 μl of media and equal volumes of DMSO for controls. For short-term treatments, a single dose of the drug or DMSO control was directly diluted in neuronal or glial maintenance media. For maturing neuronal cultures’ experiments, partial media exchanges were performed every 72 hours to maintain a sustained dCBP-1 drug activity. For recovery experiments, total media was exchanged every 72 hours to maintain a sustained dCBP-1 drug activity.

### Dendritic branching and length quantification

Hippocampal neuronal primary cultures were prepared as stated above and dCBP-1 (500nM) or DMSO were added at DIV1. Cultures were placed in an Incucyte® Live-Cell Analysis System (Sartorius) equipment and maintained until DIV24 by performing partial-media exchange every 3 days to ensure dCBP-1 activity throughout the full experimental set up. Brightfield images were acquired at 20x magnification in Incucyte® obtaining 16 images per well every hour (N=4). Neuronal dendrite length and branching parameters were measure using the *Neuronal Tracking* Incucyte® implemented plugin and data points were averaged every 12 hours for final analyses.

### CBP and p300 immunostaining and imaging analysis

Neuronal and glial mix culture from the 24-well plates were fixed at 12 minutes at room temperature in 4% paraformaldehyde, washed with PBS and permeabilized with PBS-T (PBS containing 0.2% Triton X-100). Blocking solution (3% Normal Calf Serum in PBS-T) was added for 1 hour in agitation to the samples prior overnight incubation at 4°C with primary antibodies: anti-CBP (Santa Cruz sc-7300; 1:100) and anti-p300 (Santa Cruz sc-585; 1:100). Secondary antibodies (Alexa Fluor IgG 488; Alexa Fluor IgG 594 both 1:500; Invitrogen) were incubated at room temperature for 2 hours with gentle agitation. Each sample was mounted with 10uL of DAPI fluoromount; SouthernBiotech. Image acquisition was performed using an Olympus FV1200 inverted laser-scanning confocal microscope equipped with a PLAPON 60×/1.40 NA oil-immersion objective.. The fluorescence channels were acquired sequentially, and crosstalk between fluorophores was eliminated by adjusting the detector spectral ranges and ii0 sequential scanning for image acquisition. At least five non-overlapping fields of view were acquired per sample. For each field, a z-stack spanning the full height of nuclear populations was acquired at 1 µm intervals. Laser power, detector settings, pinhole diameter, scan speed, confocal zoom, image resolution, and other acquisition parameters were kept constant between control and treatment groups within each experiment.

### CBP and p300 immunofluorescence nuclear quantifications

For data analysis, the imaging files were analyzed following a custom macro script developed for Fiji ImageJ version 1.54p software to evaluate nuclear fluorescence intensity in individual cells. Raw .oib files were imported using Bio-Formats, and z-stack maximum-intensity projections were generated for each channel. Nuclei were segmented from the DAPI channel using Huang automatic thresholding and watershed separation and stored as individual region of interest (ROI). Apart from the nuclear ROI a corresponding perinuclear ROI was generated by enlarging the nuclear ROI by a factor of 1.1 and subtracting the original nuclear area from the enlarged selection. After rolling-ball background subtraction with a radius of 90 pixels, mean fluorescence intensity was measured within the nuclear and perinuclear ROIs in each fluorescence channel. ROI-overlay images were saved and inspected to assess segmentation accuracy.

### Synaptic markers immunocytochemistry analyses

For synaptic components analysis in neuronal primary cultures, cultured hippocampal neurons were treated at DIV14 with either DMSO or dCBP-1 for 96 h or 1 week. Cells were then fixed with 4% (w/v) paraformaldehyde in phosphate-buffered saline (PBS) for 15 min at room temperature (RT). Following three washes in PBS, cells were permeabilized and blocked in blocking solution containing PBS, 10% goat serum, and 0.2% Triton X-100 for 30 min at RT. Neurons were incubated for 1 h at RT with the following primary antibodies diluted in blocking solution: mouse anti-MAP2 (1:500, Sigma-Aldrich, M9942), guinea pig anti-VGLUT1 (1:500, Synaptic Systems, #135304), mouse anti-Synapsin I (1:500, Synaptic Systems, #106011), and rabbit anti-Homer1 (1:500, Synaptic Systems, #160003). After three washes with PBS, coverslips were incubated for 1 h at RT with the appropriate secondary antibodies: Alexa Fluor 488 goat anti-mouse IgG (1:1000, Thermo Fisher Scientific #A-11001), Alexa Fluor 568 goat anti-rabbit IgG (1:500, Thermo Fisher Scientific, #A-11011), and Alexa Fluor 647 goat anti-guinea pig IgG (1:500, Thermo Fisher Scientific, #A-21450). Coverslips were subsequently washed three times with PBS and mounted using Immu-Mount (Thermo Fisher Scientific) containing 1.5 mg/ml 4′,6-diamidino-2-phenylindole (DAPI; Sigma-Aldrich) for nuclear staining. Samples were imaged using a Zeiss LSM880 laser-scanning confocal microscope equipped with a 63× oil-immersion objective. Acquisition parameters were kept constant between control and treated groups within each experiment. Fluorescence intensities were quantified using ImageJ software and normalized to the corresponding control condition. Data were obtained from 1–3 independent experiments. Normalized data were analyzed using two-tailed one-sample *t*-tests against a theoretical mean of 1.

### RNA extraction

Total RNA from neuronal and glial cultures was extracted using TRI Reagent® by mechanical dissociation using a p1000 pipette to detach and homogenize cell lysates. RNA was extracted as previously described (Lipinski et al., 2020) by using first Phenol:Chloroform:Isoamyl Alcohol (25:24:1) and two consecutive rounds of Chloroform before a precipitation step with isopropanol and RNAse-free glycogen. Genomic DNA was eliminated with DNAse T treatment and DNA-free RNA was again repurified using Phenol / Chloroform extraction and precipitation. Finally, samples were diluted in 20-25 µl of DEPC water previously heat at 65 °C and stored at −80° C and RNA concentration was evaluated using NanoDrop™ One.

### RNA-seq: Library processing and sequencing

Libraries were prepared using the TruSeq Stranded Total RNA Library Prep kit (ref. 20020597 for Human/Mouse/Rat) according to the manufacturer’s protocol, to convert total RNA into a library of template molecules of known strand origin and suitable for subsequent cluster generation and DNA sequencing. Briefly, 300 ng of total RNA were used to remove ribosomal RNA (rRNA) using biotinylated, target-specific oligos combined with Ribo-Zero rRNA removal beads. An extra step using Sera-Mag beads (ref. 30152103010150, GE Healthcare) was added to the Illumina protocol to improve rRNA depletion. After the rRNA was depleted, the remaining RNA was purified with RNAClean XP beads (ref. A63987, Beckman Coulter). Following purification, the RNA was fragmented using divalent cations under elevated temperature and primed with random hexamers for cDNA synthesis, that was performed using reverse transcriptase (SuperScript II; ref. 18064-014, Invitrogen). Then, second strand cDNA was synthesized, incorporating dUTP in place of dTTP and generating blunt-ended ds cDNA. A single ‘A’ nucleotide was added to the 3’ ends of the blunt fragments (A-tailing) and immediately afterwards the Truseq adapter was ligated. Finally, PCR selectively enriched those DNA fragments that had adapter molecules on both ends. The PCR was performed using Unique Dual Indexes and the master mix provided with the kit. All purification steps during library preparation were performed using AgenCourt AMPure XP beads (ref. A63882, Beckman Coulter). Final libraries were analyzed using Bioanalyzer DNA 1000 or Fragment Analyzer Standard Sensitivity (ref. 5067-1504 or ref. DNF-473, Agilent) to estimate the quantity and validate the size distribution and were then quantified by qPCR using the KAPA Library Quantification Kit KK4835 (ref. 07960204001, Roche). Libraries were sequenced 1 * 51 bp on Illumina’s NextSeq2000.

### RNA- seq data analysis

For RNA-seq data analysis, Trim Galore (v0.6.4_dev) was used for adapter trimming, and STAR (v2.6.1) (Dobin et al., 2013) was used for alignment to the mm10 mouse genome. Files were filtered using −q 30 in Samtools (v1.7) (Li et al., 2009) . Gene counts were obtained using Rsubread (v2.20) with the Ensembl GTF annotation (version 99) in reverse-stranded mode. For the calculations regarding exons, introns and intergenic reads, exons were calculated as mentioned above. For introns “Gene” was used to calculate all the read in the genes, then “Gene” – “Exons” was used to obtain reads associated to Introns. Intergenic reads were calculated by the quantity of aligned reads – those reads in “Gene”. Gene annotation was done by using the GTF file or Biomart(v2.26.1) (Durinck et al., 2005) when needed. For the PCA in Astrocytes vs Neurons, limma (v.3.62.2) (Ritchie et al., 2015) was used to remove the effects done to “Tissue” then the top 5000 genes were used to perform the PCA. Differential expression analysis was performed with DESeq2 (v1.46) (Love et al., 2014) using the LRT for the time-course, and Wald test was used for the individual comparisons. BigWig files were generated using deepTools (v3.5.6) (Ramírez et al., 2016) with RPKM normalization. Heatmaps and K-means clustering were performed using ClusterGVis (Zhang 张俊 et al., 2026) in combination with ClusterProfiler (Yu et al., 2012) and EnrichR database (Kuleshov et al., 2016) for Gene Ontology analyses. ChIP-seq data metaprofiles were generated with DeepTools. Synaptic Gene Ontology analyses were performed using SynGo (Koopmans et al., 2019).

In RNA-seq data from glial dCBP-1 Recovery experiments fully recovered genes were defined as those presenting a significantly higher expression in the Recovery condition compared to samples treated with dCBP-1 for two weeks and not significant compared to samples treated with DMSO for two weeks); partially recovered genes as those presenting a significantly higher expression in the Recovery condition compared to samples treated with dCBP-1 for two weeks but still presenting a significantly lower expression compared to samples treated with DMSO for two weeks) and not recovered genes as those presenting a significantly lower expression in the Recovery condition compared to samples treated with DMSO for two weeks and not significantly higher than samples treated with dCBP-1 for two weeks).

### Electrophysiological recordings

Patch-clamp recordings were obtained at room temperature from cultured hippocampal neurons at DIV 14. Cells were visualized by using infrared light and differential interference contrast (DIC) microscopy through a 40x Zeiss water-immersion objective using an upright microscope (Axioskop FS, Zeiss). During experiments, cultured neurons were continuously perfused with fresh extracellular solution containing (in mM): 130 NaCl, 2.7 KCl, 10 HEPES, 2 MgCl_2_, 10 Glucose, 2 CaCl_2_ (pH 7.4). Cells were whole-cell voltage-clamped at a holding potential of −60 mV using an EPSC10 amplifier controlled by Patchmaster software (HEKA). Sampling intervals and low-pass filter settings were 20 µs and 2.9 kHz, respectively. Patch-pipettes were pulled from borosilicate glass capillaries with filament (World Precision Instruments) on a P-97 Flaming/Brown Puller (Sutter Instruments) to have an open-tip resistance of 3-5 MW when filled with a pipette solution containing (in mM) 120 KGluconate, 20 KCl, 10 EGTA, 2 MgCl_2_, 2 Na_2_ATP, 10 HEPES (∼315 mOsmol/l, pH 7.3). Presynaptic inputs were stimulated with a glass electrode filled with extracellular solution and placed on a nearby cultured neuron ≥80 μm away from the patched cell. Postsynaptic currents (ePSCs) were evoked by applying brief electrical pulses (140 μs, 10–50 V). Stimulus intensity was adjusted to avoid polysynaptic components. Spontaneously occurring EPSCs (sEPSCs) were recorded in continuous current sweeps in the absence of TTX. Action potentials (APs) were elicited in current-clamp recordings with the injection of 500 ms square current pulses growing in their amplitudes from 0 to 450 pA in steps of 50 pA. All offline analysis of electrophysiological data was performed in Igor Pro (Wavemetrics). Voltage-clamp recordings were leak-corrected and low-pass filtered with a cut-off frequency fc = 3 kHz using a 10-pole digital Bessel filter. sPSCs were detected by applying a sliding template-based detection algorithm.

### Phagocytosis Assay

Primary glial cells were seeded in 48-well plates at a density of 40,000 cells/well. Upon reaching maturity, phagocytic activity was assessed using pHrodo™ Red *Escherichia coli* BioParticles™ conjugate (Thermo Fisher Scientific, P35361). The BioParticles were reconstituted in a pH-neutral saline solution to a 1 mg/mL stock concentration and sonicated for 10 minutes to ensure uniform suspension. The particles were added to the cells to reach a final concentration of 100 μg/mL in a total volume of 100 μl per well. Plates were immediately transferred to an IncuCyte® Live-Cell Analysis System (Sartorius) and maintained at 37°C. Images were captured every 30 minutes for 15 hours using a 20x objective. To quantify phagocytosis, red fluorescence metrics—including average and total integrated intensity and red object count per image—were retrieved using the IncuCyte® software.

### Wound-Healing (Scratch) Assay

To evaluate glial cell migration and physiological recovery *in vitro*, a wound-healing assay was performed on mature mixed glial cultures. Cells were grown in ImageLock 96-well plates (Sartorius, BA-04855) and a precise mechanical scratch was manually created in the cell monolayer using a sterile 10 μl pipette tip. Following the scratch, the culture medium was exchanged twice to remove dislodged cells and prevent unwanted reattachment. The plates were then placed in the IncuCyte® system at 37°C. Automated imaging was performed using a 4x objective at 1-hour intervals for a total duration of 120 hours (5 days). Migration was quantified by calculating the Relative Wound Density (RWD) via the IncuCyte® integrated software.

### Statistical analysis

All statistical analyses were performed using GraphPad Prism (v11.1.0; GraphPad Software, La Jolla, CA, USA) and RStudio v3.5.1. Every set of data was assessed for normality using the GraphPad implemented Saphiro-Wilk test. Two-tailed unpaired Student’s t-test was applied for comparisons between two independent groups when following a normal distribution. When more than two groups were compared, a one-way or two-way analysis of variance (ANOVA) was performed according to the experimental groups, followed by a Tukey’s or Bonferroni’s multiple comparison test as post hoc test. Gene expression values were normalized as row-wise Z-scores, and differences across conditions were evaluated using the Kruskal-Wallis test, with post hoc pairwise Wilcoxon rank-sum tests corrected for multiple comparisons using the Dunn’s Test. Statistical significance was set at p < 0.05.

## Data availability

The genomic datasets generated in this study will be deposited in the Gene Expression Omnibus (GEO) repository upon acceptance of the manuscript.

## Code availability

Code used for RNA-seq data analysis and imaging analysis automatization will be made available under demand.

## SUPPLEMENTARY FIGURE LEGENDS

**Supplementary Figure 1 related to Figure 1. dCBP-1 effect in maturing neuronal cultures and neuronal RNA-seq time-course QC. a.** Schematic representation of dCBP-1 experiment assessing the impact of KAT3 elimination during neuronal primary cultures maturation using Incucyte®. Created in Biorender. Fontana, T.(2024). **b.** Neurite length quantification (mm/ mm^2^) along 24 days time window of neuronal primary cultures treated with DMSO (black) or dCBP-1 (red). Data is represented as mean ± SEM. \*\*\**P* < 0.001 as compared with DMSO using Two-way Anova. **c.** Branching points quantification (branching points / mm^2^) along 24 days time window of neuronal primary cultures treated with DMSO (black) or dCBP-1 (red). Data is represented as mean ± SEM. \*\*\**P* < 0.001 as compared with DMSO using Two-way Anova. **d.** Representative images of brightfield Incucyte® acquisitions of DMSO (top) and dCBP-1 (bottom) treated neuronal cultures at DIV1, DIV3, DIV6, DIV12 and DIV24. Scale bar, 200 µm. **e.** Summary table of RNA-seq mapped reads per replicate along the dCBP-1 neuronal primary cultures time course experiment. **f.** Bar graphs showing the % of reads mapping at different genomic regions (exons, introns and intergenic) in the neuronal dCBP-1 time-course RNA-seq data. *P< 0.05, \*\**P* < 0.01 as compared with DMSO using Two-way Anova and Tukey’s multiple comparison test. **g.** Heatmap showing CBP and p300 normalized gene expression in DMSO and dCBP-1 (3, 6, 24 and 48 hours) treated neurons from RNA-seq time-course.

**Supplementary Figure 2 related to Figure 2. Additional dCBP-1 neuronal time-course RNA-seq data GO analyses. a.** Synaptic GO (SynGO) analysis of down-regulated genes in dCBP-1 time-course RNA-seq data at 3, 6, 24, and 48 hours of dCBP-1 treatment. **b.** Synaptic GO (SynGO) analysis of up-regulated genes in dCBP-1 time-course RNA-seq data at 3, 6, 24, and 48 hours of dCBP-1 treatment. **c.** Gene Ontology SynGO bubble plots showing enriched GO terms associated to the genes belonging to the different clusters generated in Figure 2d. Bubble color indicates statistical significance (AdjPvalue) and size the gene counts associated to the term. **d.** Gene Ontology ChIP-X Enrichment Analysis (CheA) bubble plots showing enriched GO terms associated to the genes belonging to the different clusters generated in Figure 2d. Bubble color indicates statistical significance (AdjPvalue) and size the gene counts associated to the term.

**Supplementary Figure 3 related to Figure 3. KAT3 depletion by dCBP-1 induces a morphological and functional immature phenotype in cultured hippocampal neurons characterized by reduction of synapse number and functional enhanced spontaneous neurotransmission. a.** Representative confocal images of DIV14 hippocampal neurons treated with DMSO (control) or dCBP-1 for 96h or 1 week, followed by immunostaining for Synapsin 1 (green). Nuclei were counterstained with DAPI (blue). Scale bars, 10 μm. **b.** Quantification of the number of synapsin 1-positive puncta per image (left) and quantification of fluorescence intensity (right), both normalized to DMSO-treated neurons (set to 1). Violin plots represent the distribution of values, with dashed lines indicating the median and quartiles. Quantifications were based on 24 and 17 images in the DMSO and dCBP-1 1-week groups, respectively, collected across 1–3 independent experiments. **c.** Representative traces of spontaneously occurring PSCs (sPSCs) recorded from mass-cultured hippocampal neurons treated with DMSO (control, salmon) or dCBP-1 for 96 h (violet). Right traces show corresponding mean spontaneous PSC waveforms. **d.** Scatter plots and bar graphs showing individual and mean values respectively, for frequency of spontaneous PSCs in neurons treated with DMSO (control), or dCBP-1 for 24h, 48h, and 96h. Statistical significance in **a-b** was determined using two-tailed one-sample t-tests against a theoretical value of 1. In **d**, error bars indicate SEM. Statistical comparisons were performed using one-way ANOVA followed by Tukey test. **P < 0.01.

**Supplementary Figure 4 related to Figure 4. Additional dCBP-1 glial time-course RNA-seq data analyses. a.** Summary table of RNA-seq mapped reads per replicate along the dCBP-1 glial primary cultures time-course experiment. **b.** Heatmap showing CBP and p300 normalized gene expression in DMSO and dCBP-1 (6, 24 and 48 hours) treated glial cultures from RNA-seq time-course. **c.** Bar graphs showing the % of reads mapping at different genomic regions (exons, introns and intergenic) in the glial dCBP-1 time-course RNA-seq data. \*\**P* < 0.01 as compared with DMSO using Two-way Anova and Tukey’s multiple comparison test. **d.** Gene body metaprofiles of mouse hippocampal CBP, p300 and H3K27ac ChIP-seq (Lipinski et al., 2020) signal enrichment in genes down-(blue) and up-regulated (red) in glial primary cultures treated with dCBP-1.

**Supplementary Figure 5 related to Figure 5. Additional dCBP-1 glial time-course RNA-seq data GO and functional analyses. a.** Synaptic GO (SynGO) analysis of down-regulated genes in dCBP-1 time-course RNA-seq data at 6, 24, and 48 hours of dCBP-1 treatment. **b.** Synaptic GO (SynGO) analysis of up-regulated genes in dCBP-1 time-course RNA-seq data at 6, 24, and 48 hours of dCBP-1 treatment. **c.** IGV genome tracks of dCBP-1 time-course RNA-seq data in glial primary cultures showing the progressive downregulation at the glial *Gfap*, *Olig1* and *Cx3cr1* gene loci. **d.** Gene Ontology ChIP-X Enrichment Analysis (CheA) bubble plots showing enriched GO terms associated to the genes belonging to the different clusters generated in Figure 5a. Bubble color indicates statistical significance (AdjPvalue) and size the gene counts associated to the term. Arrows indicate interesting regulators for each cluster due to the common cellular model or to the direct link with KAT3 proteins. **e.** Red Object Count quantification of pHrodo phagocytosis shown per hour or with all acquisition points together. Data is shown as mean ± SEM. *ns, non-significant, ****P<0,0001 as compared with DMSO using Two-way Anova and Tukey’s multiple comparison test for mean group effect. **f.** pHrodo signal quantification as Average Red Object Integrated Intensity (RCU x µm^2^) shown per hour or with all acquisition points together. Data is shown as mean ± SEM. **P<0,01, ****P<0,0001 as compared with DMSO using Two-way Anova and Tukey’s multiple comparison test for mean group effect.

**Supplementary Figure 6 related to Figure 6. Additional analyses of dCBP-1 effect in glial and neuronal identity programs. a.** Dot blot showing hippocampal cell identity markers normalized mean expression in neuronal (purple) or glial (green) RNA-seq time-course data in DMSO or dCBP-1 (6, 24 and 48 hours) treated cells. **b.** Gene Ontology ChIP-X Enrichment Analysis (CheA) bubble plots showing enriched GO terms associated to the genes belonging to the different gene identity clusters generated in Figure 6e. Bubble color indicates statistical significance (AdjPvalue) and size the gene counts associated to the term. **c.** Gene body metaprofiles of mouse hippocampal CBP, p300 and H3K27ac ChIP-seq (Lipinski et al., 2020) signal enrichment in neuronal (top, clusters 2, 3 and 4) and glial (bottom, clusters 1, 5 and 6) identity clusters originated in Figure 6e. **d-e.** Area fill graph showing the normalized gene expression as % to DMSO of developmental cortical modules (Gao et al., 2025) in neuronal (d) and glial (e) RNA-seq time course data for DMSO and dCBP-1 (6, 24 and 48 hours) treated cells.

**Supplementary Figure 7 related to Figure 7. Complementary analyses of dCBP-1 washout experiment in glial cells. a.** Representative confocal images of DAPI (cyan), CBP (green) and p300 (red) in glial primary cultures treated for 6, 24, 48, 72h and 96 hours with a single administration of dCBP-1 or 96 hours with DMSO. Scale bar, 20 µm. **b.** Violin plots showing CBP and p300 nuclear integrated density as % to DMSO controls in DMSO and dCBP-1 (6, 24, 48, 72 and 96 hours) treated cells (N=4). ****P<0,0001 as compared with DMSO (Mann-Whitney test). **c.** On the top, representative confocal images of DAPI (cyan), CBP (green) and p300 (red) in glial primary cultures treated for 1 week with DMSO (DMSO 1W) or dCBP-1 (dCBP-1 1W). Scale bar, 20 µm. On the bottom, violin plots showing CBP and p300 nuclear integrated density as % to DMSO controls (DMSO 2W) in DMSO 2W, dCBP-1 2W and Recovery conditions (N=4). ns, non-significant, ****P<0,0001 as compared with DMSO 2W (Mann-Whitney test). **d.** Summary table of RNA-seq mapped reads per replicate along the dCBP-1 glial primary culture recovery experiment. **e.** Spearman correlation heatmap of RNA-seq samples from glial dCBP-1 recovery experiment. **f.** Bar graphs showing the % of reads mapping at different genomic regions (exons, introns and intergenic) in the glial dCBP-1 recovery RNA-seq data. No significant differences were identified using Two-way Anova and Tukey’s multiple comparison test.

**Supplementary Figure 8 related to Figure 8. Complementary analyses of dCBP-1 washout RNA-seq and functional experiment in glial cells. a.** Bar graphs showing the normalized expression of fully recovered (left), partially recovered (mid) and not recovered (right) genes in RNA-seq data from DMSO 2W (red), dCBP-1 2W (green) and Recovery (blue) samples. Data is shown as mean ± SEM. **b.** Area fill graph showing the % of genes belonging to the different developmental cortical modules (Gao et al., 2025) found in the distinct glial RNA-seq recovery categories described in Figure 8a. In blue, fully recovered genes; black, partially recovered genes and red, not recovered genes. **c.** Gene Ontology ChIP-X Enrichment Analysis (CheA) bubble plots showing enriched GO terms associated to the gene recovery categories described in Figure 8a (fully recovered, partially recovered and not recovered). Bubble color indicates statistical significance (AdjPvalue) and size the gene counts associated to the term. **d.** Stacked bar graph showing the % of glial identity genes comprised among the lists of fully recovered, partially recovered and not recover genes according to Figure 8a. **e.** IGV genome tracks of dCBP-1 recovery RNA-seq data in glial primary cultures showing the expression of the glial *Gfap*, *Olig1* and *Cx3cr1* glial markers. **f.** Red Object Count quantification of pHrodo phagocytosis shown per hour or with all acquisition points together in dCBP-1 recovery experiments. Data is shown as mean ± SEM. *ns, non-significant, ****P<0,0001 as compared with DMSO using Two-way Anova and Tukey’s multiple comparison test for mean group effect. **G.** pHrodo signal quantification as Average Red Object Integrated Intensity (RCU x µm^2^) shown per hour or with all acquisition points together in dCBP-1 recovery experiments. Data is shown as mean ± SEM. ****P<0,0001 as compared with DMSO using Two-way Anova and Tukey’s multiple comparison test for mean group effect.

