## Supplementary Figures for "Dynamic Modulation of Neural Cell Identity Maintenance via Lysine Acetyltransferase Type 3 Protein Targeted Degradation"

### Supplementary Figure 1

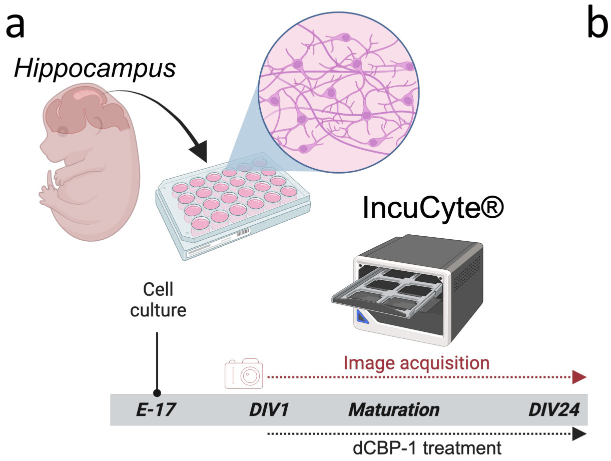


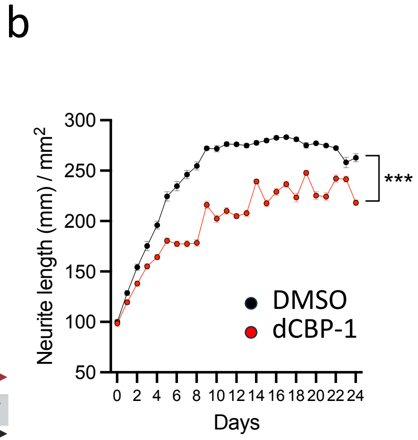


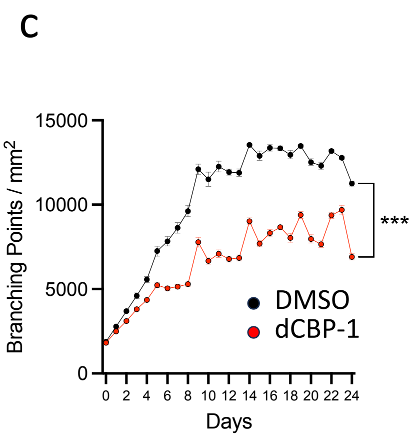


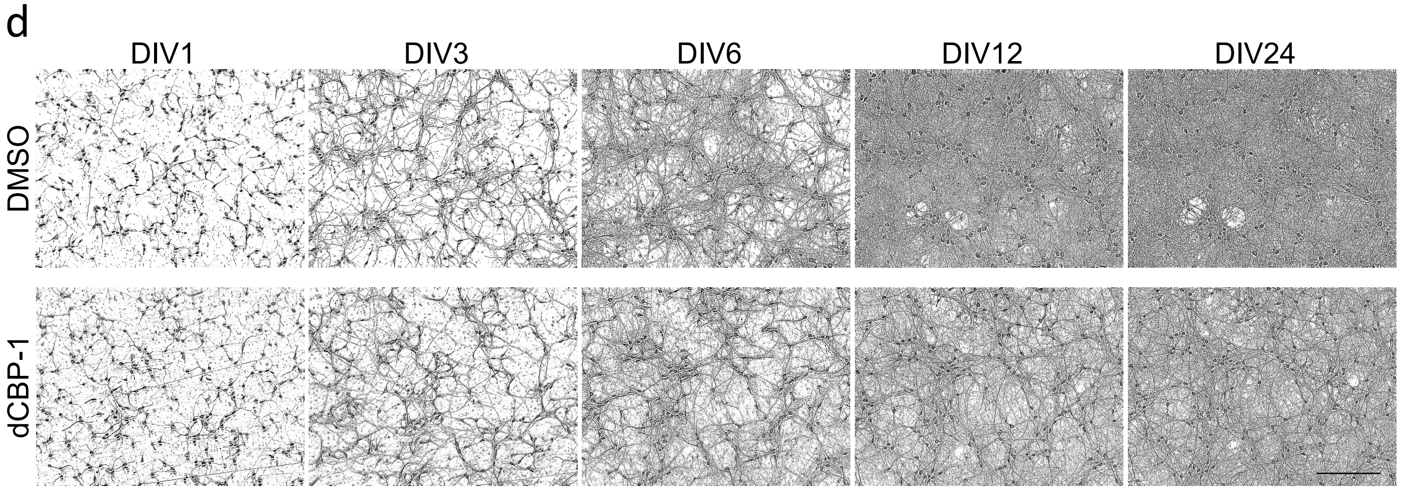


e

| Sample | Mapped reads |
|---|---|
| DMSO Rep1 | 47,145,698 |
| DMSO Rep2 | 53,979,167 |
| DMSO Rep3 | 43,862,979 |
| dCBP-1 3h Rep1 | 61,531,116 |
| dCBP-1 3h Rep2 | 59,655,167 |
| dCBP-1 3h Rep3 | 48,004,231 |
| dCBP-1 6h Rep1 | 66,558,661 |
| dCBP-1 6h Rep2 | 62,572,945 |
| dCBP-1 6h Rep3 | 57,161,005 |
| dCBP-1 24h Rep1 | 72,051,804 |
| dCBP-1 24h Rep2 | 66,119,383 |
| dCBP-1 24h Rep3 | 72,915,638 |
| dCBP-1 48h Rep1 | 54,101,182 |
| dCBP-1 48h Rep2 | 61,218,940 |
| dCBP-1 48h Rep3 | 64,797,195 |

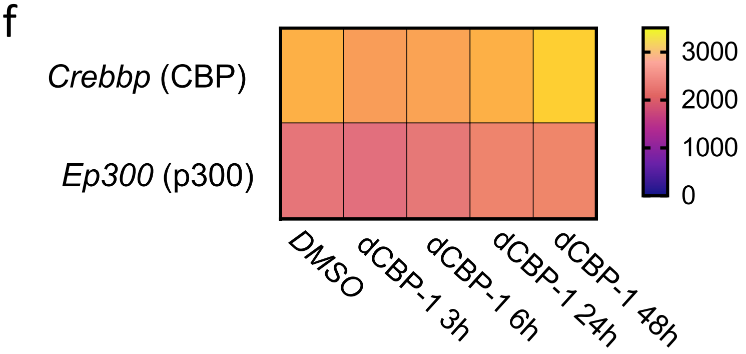


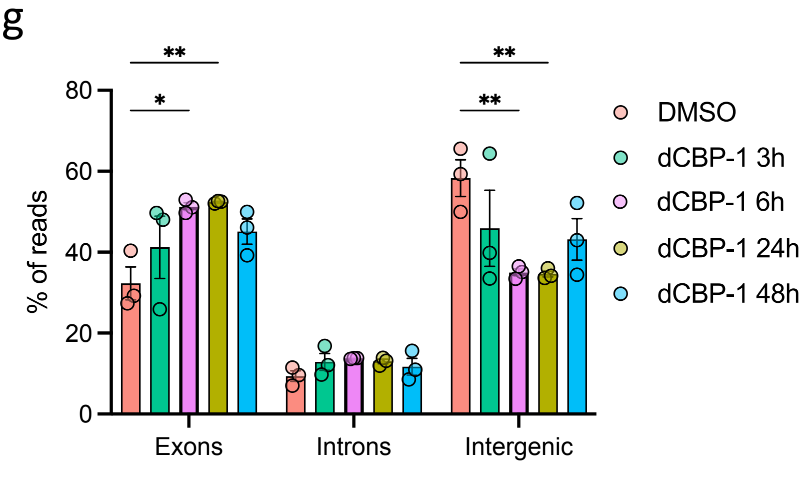

### Supplementary Figure 2

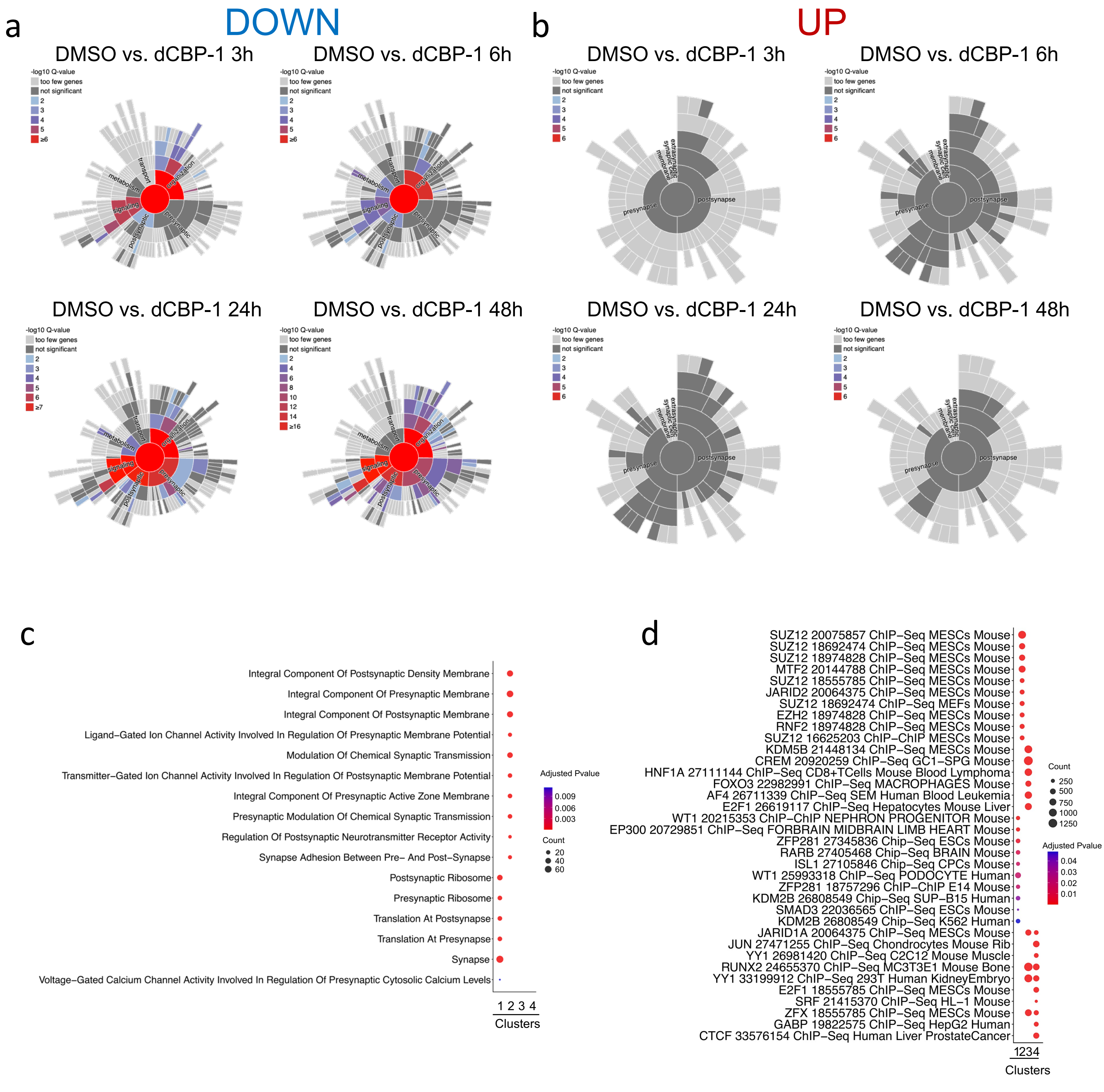

### Supplementary Figure 3

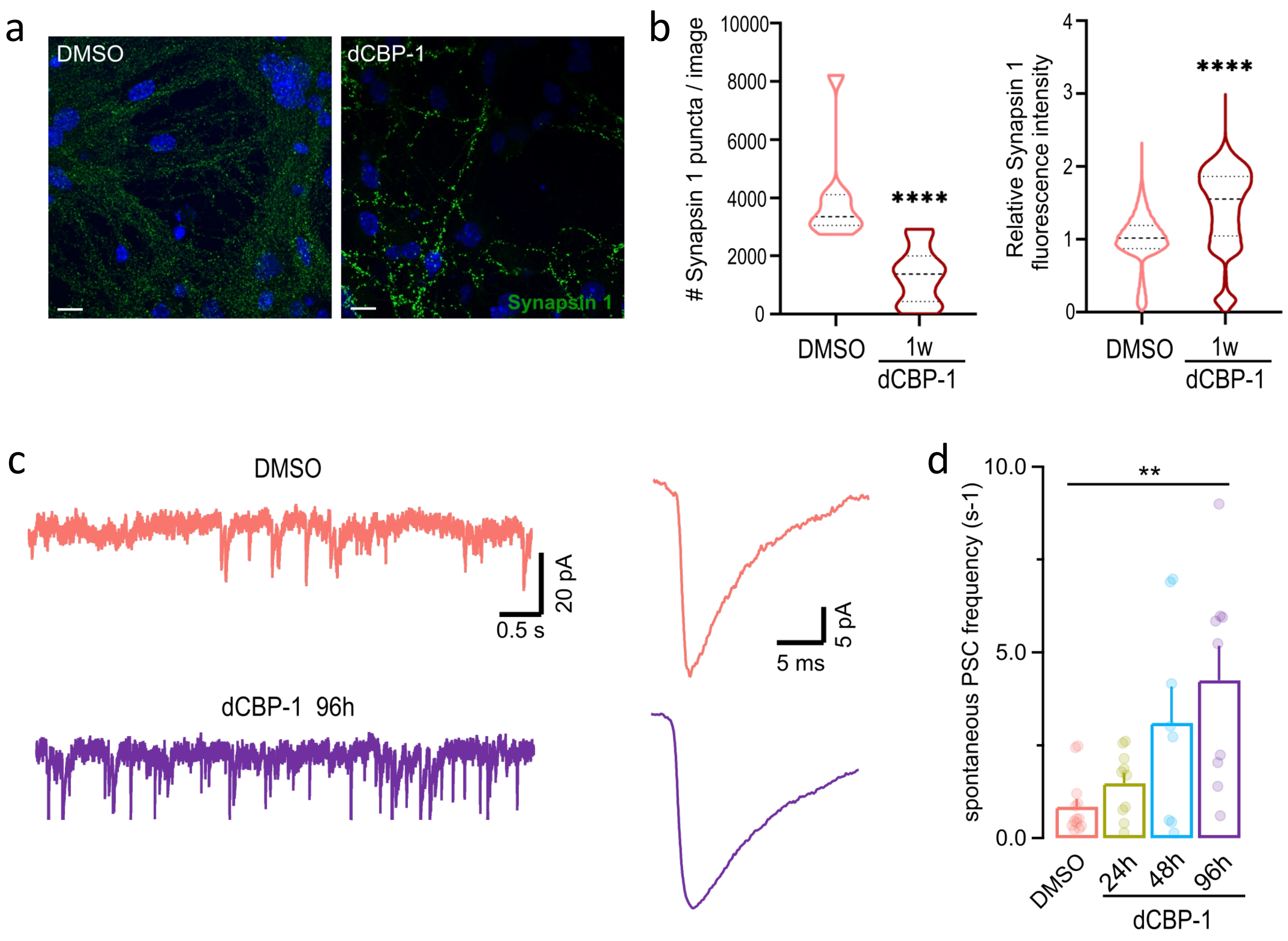

### Supplementary Figure 4

a

| Sample | Mapped reads |
|---|---|
| DMSO Rep1 | 61,752,419 |
| DMSO Rep2 | 61,731,569 |
| DMSO Rep3 | 61,127,371 |
| DMSO Rep4 | 51,387,510 |
| dCBP-1 6h Rep1 | 55,633,250 |
| dCBP-1 6h Rep2 | 62,180,717 |
| dCBP-1 6h Rep3 | 59,876,624 |
| dCBP-1 6h Rep4 | 66,161,119 |
| dCBP-1 24h Rep1 | 59,165,962 |
| dCBP-1 24h Rep2 | 57,794,779 |
| dCBP-1 24h Rep3 | 55,394,894 |
| dCBP-1 24h Rep4 | 50,490,853 |
| dCBP-1 48h Rep1 | 53,340,245 |
| dCBP-1 48h Rep2 | 49,138,336 |
| dCBP-1 48h Rep3 | 53,421,116 |
| dCBP-1 48h Rep4 | 53,736,765 |

b

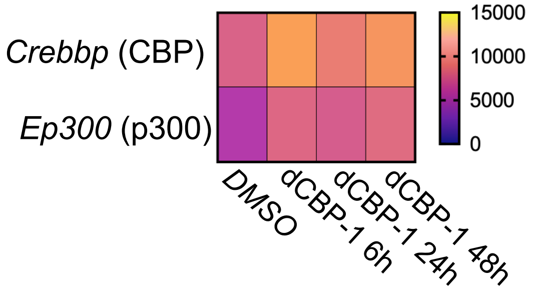


c

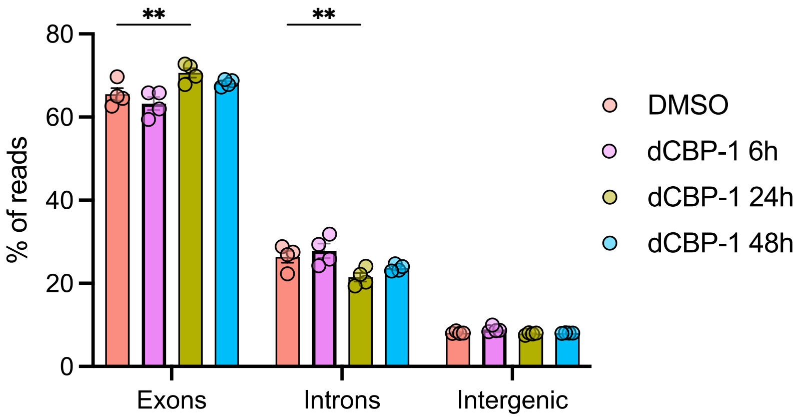


d

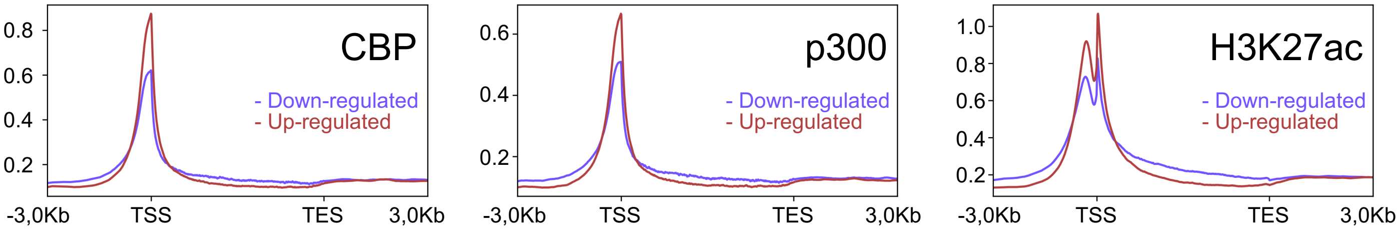

### Supplementary Figure 5

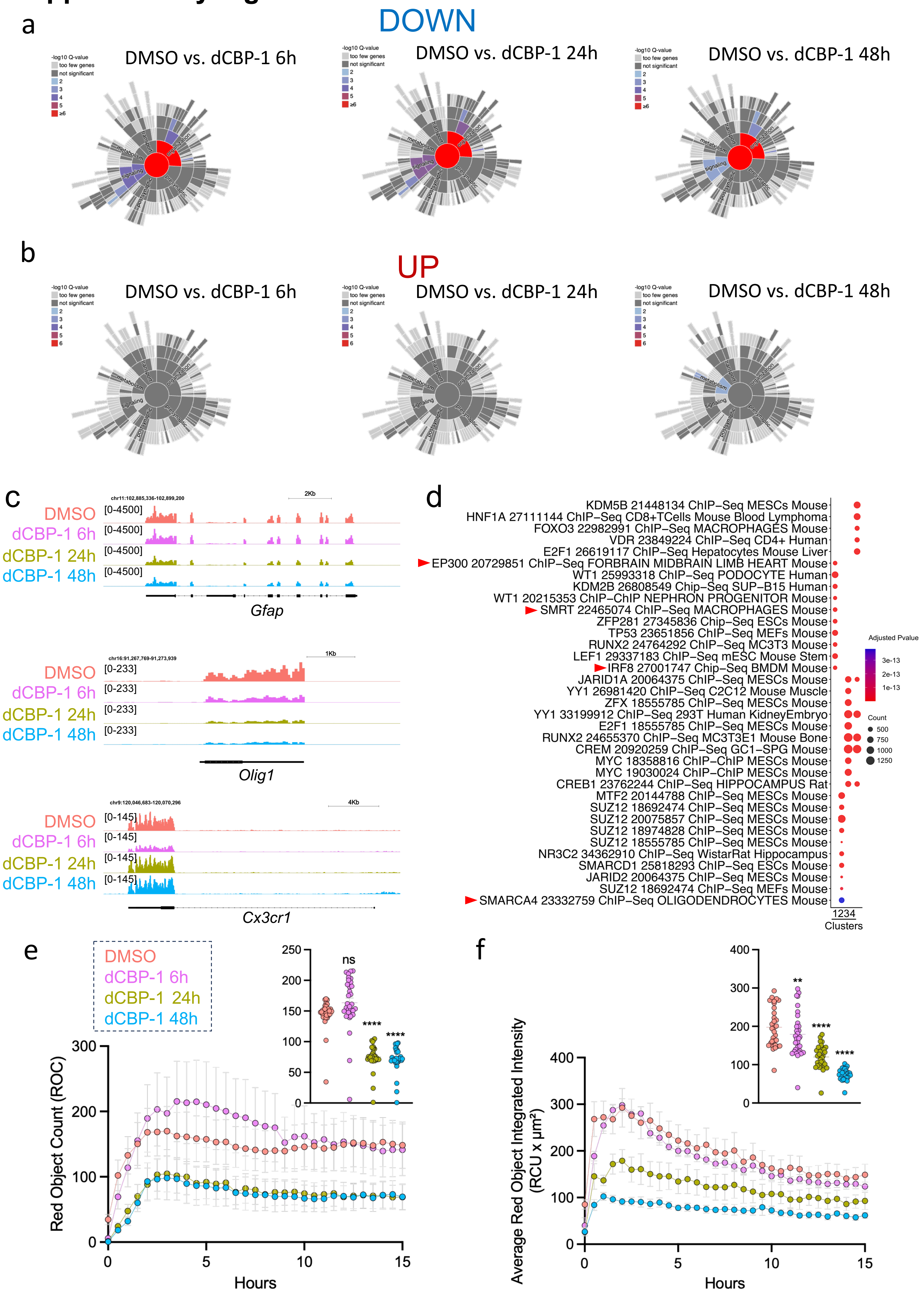

### Supplementary Figure 6

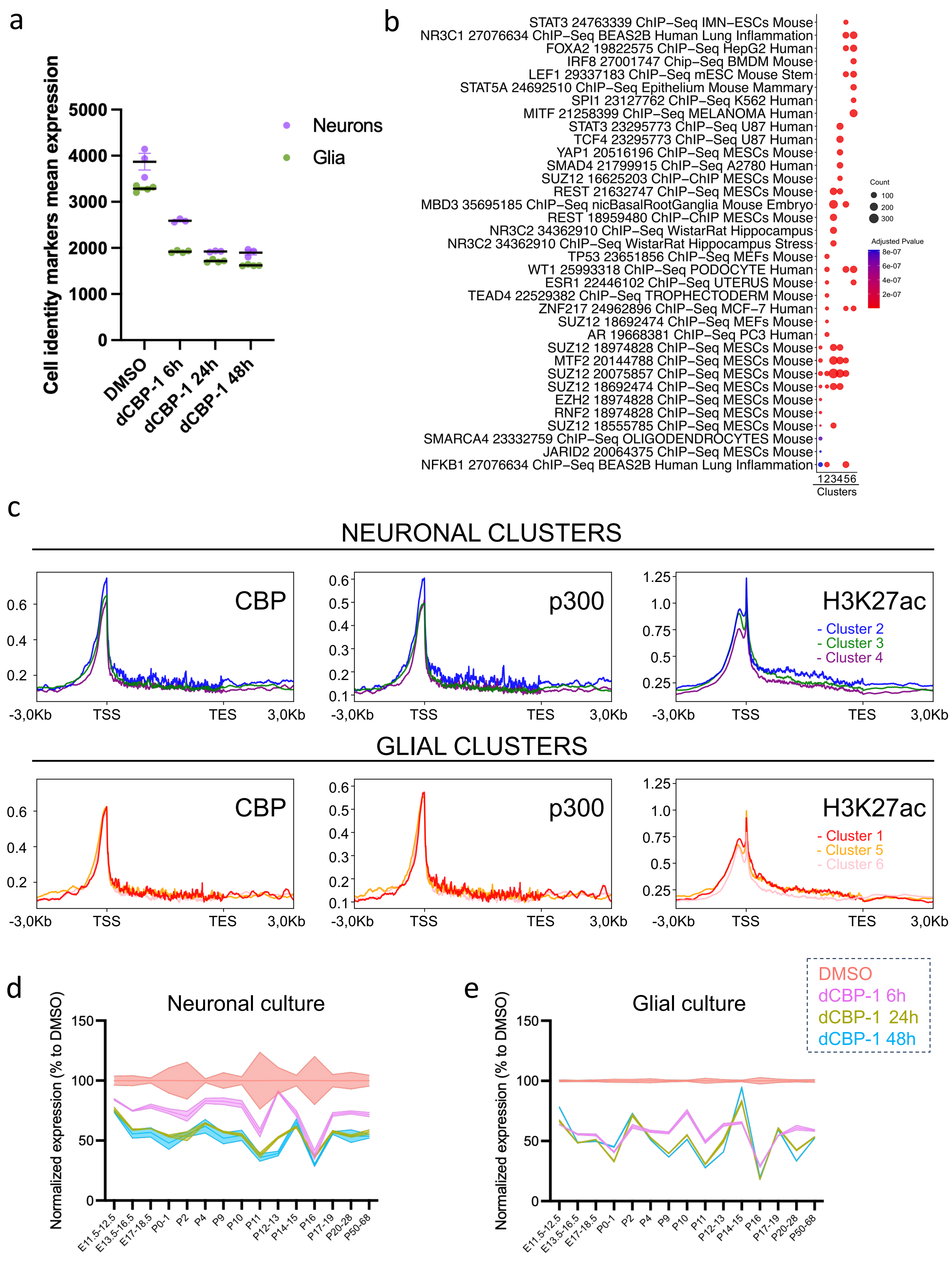

### Supplementary Figure 7

a

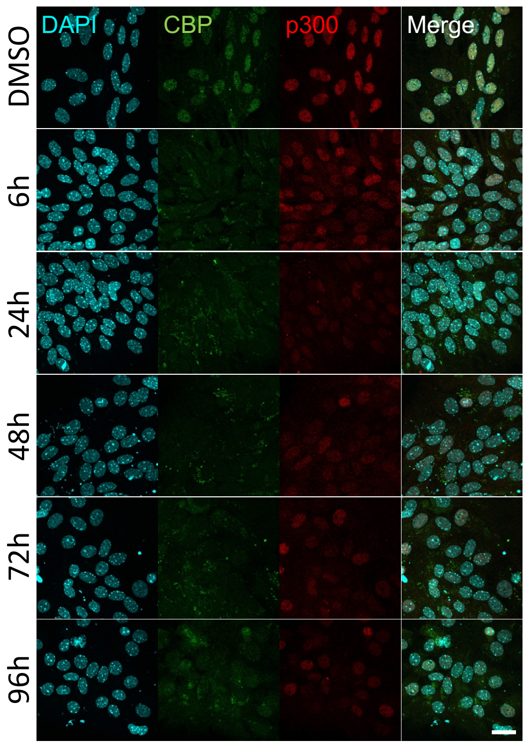


b

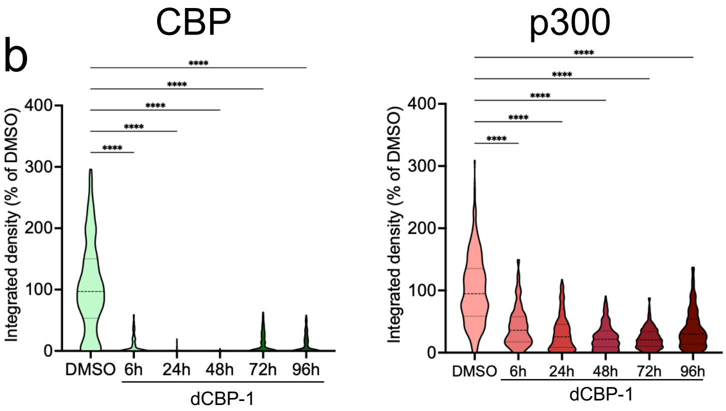


c

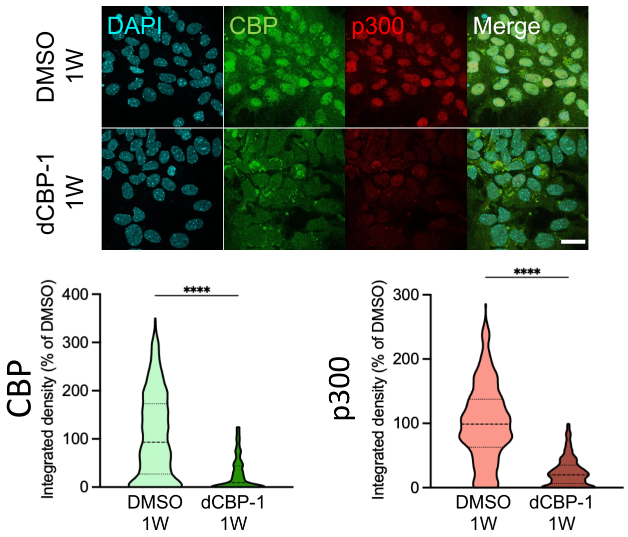


d

| Sample | Mapped reads |
|---|---|
| DMSO 1W Rep1 | 42,266,999 |
| DMSO 1W Rep2 | 46,167,673 |
| DMSO 1W Rep3 | 39,736,942 |
| DMSO 1W Rep4 | 40,980,684 |
| dCBP-1 1W Rep1 | 37,210,575 |
| dCBP-1 1W Rep2 | 42,107,973 |
| dCBP-1 1W Rep3 | 43,051,567 |
| dCBP-1 1W Rep4 | 38,698,549 |
| DMSO 2W Rep1 | 44,295,827 |
| DMSO 2W Rep2 | 43,258,235 |
| DMSO 2W Rep3 | 42,964,952 |
| DMSO 2W Rep4 | 41,797,345 |
| dCBP-1 2W Rep1 | 42,592,579 |
| dCBP-1 2W Rep2 | 43,912,877 |
| dCBP-1 2W Rep3 | 46,954,866 |
| dCBP-1 2W Rep4 | 43,925,285 |
| dCBP-1 Re Rep1 | 42,844,906 |
| dCBP-1 Re Rep2 | 43,250,782 |
| dCBP-1 Re Rep3 | 41,783,563 |
| dCBP-1 Re Rep4 | 40,630,667 |

e

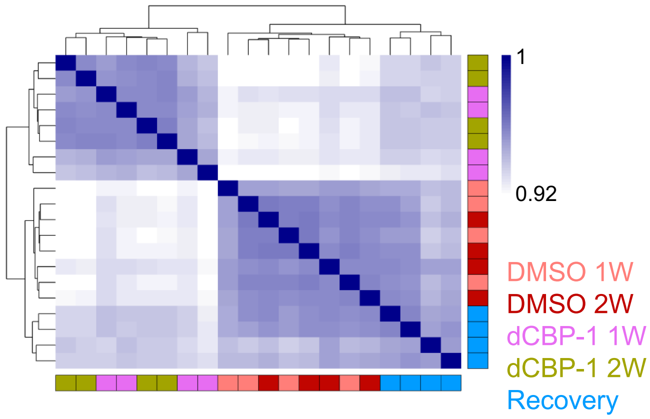


f

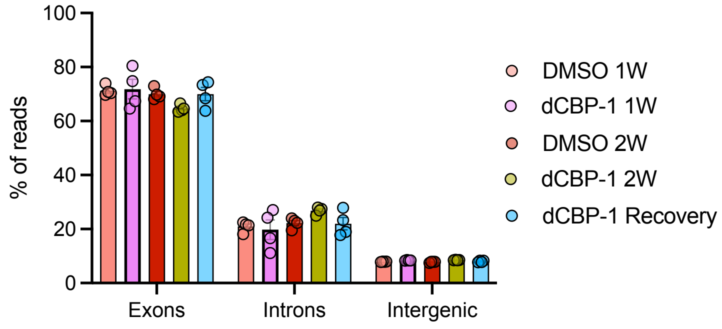

### Supplementary Figure 8

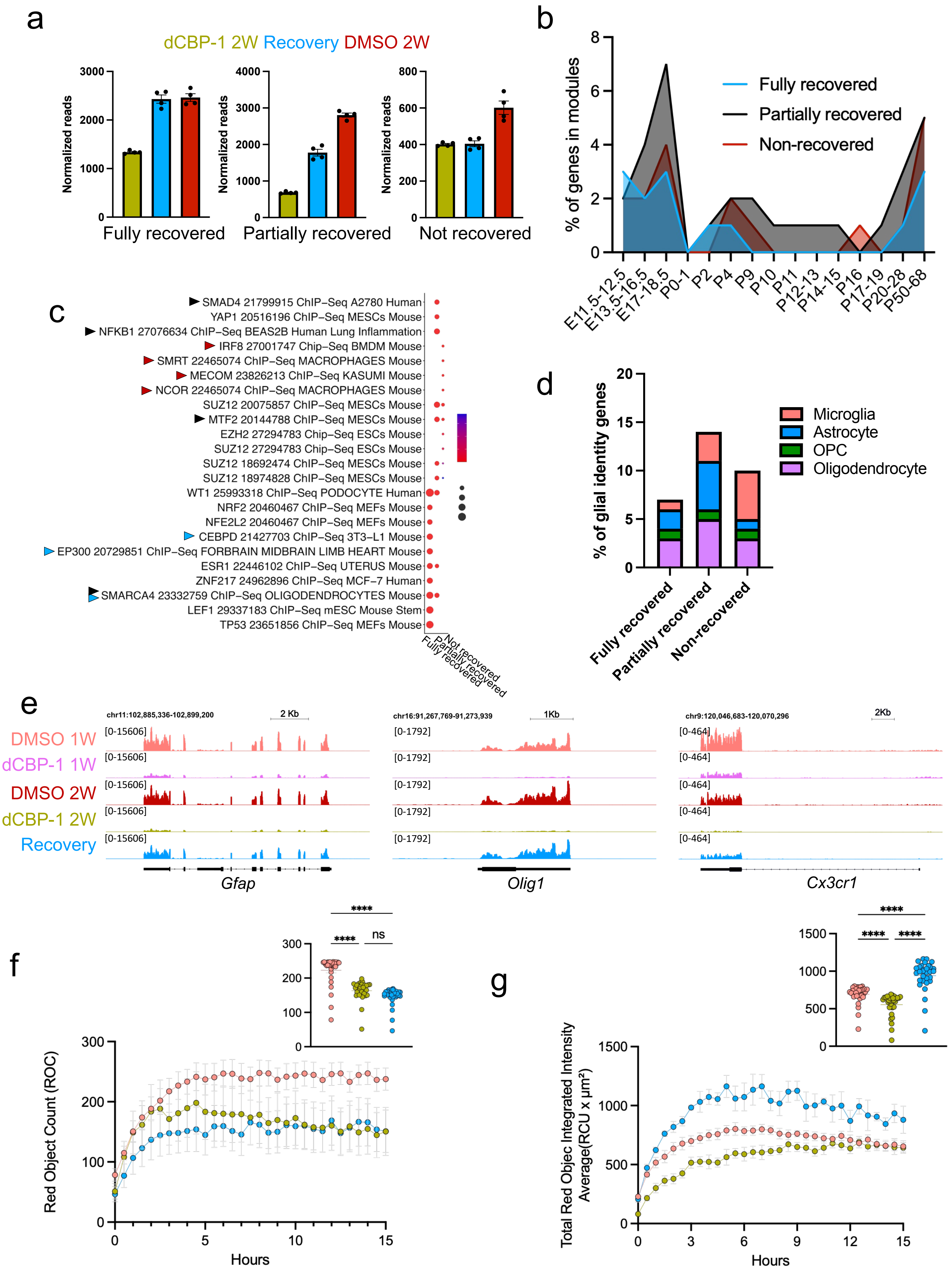
